# The neural circuitry underlying the “rhythm effect” in stuttering

**DOI:** 10.1101/2020.10.27.350975

**Authors:** Saul A. Frankford, Elizabeth S. Heller Murray, Matthew Masapollo, Shanqing Cai, Jason A. Tourville, Alfonso Nieto-Castañón, Frank H. Guenther

## Abstract

**Purpose:** Stuttering is characterized by intermittent speech disfluencies which are dramatically reduced when speakers synchronize their speech with a steady beat. The goal of this study was to characterize the neural underpinnings of this phenomenon using functional magnetic resonance imaging.

**Method:** Data were collected from 17 adults who stutter and 17 adults who do not stutter while they read sentences aloud either in a normal, self-paced fashion or paced by the beat of a series of isochronous tones (“rhythmic”). Task activation and task-based functional connectivity analyses were carried out to compare neural responses between speaking conditions and groups.

**Results:** Adults who stutter produced fewer disfluent trials in the rhythmic condition than in the normal condition. While adults who do not stutter had greater activation in the rhythmic condition compared to the normal condition in regions associated with speech planning, auditory feedback control, and timing perception, adults who stutter did not have any significant changes. However, adults who stutter demonstrated increased functional connectivity between bilateral inferior cerebellum and bilateral orbitofrontal cortex as well as increased connectivity among cerebellar regions during rhythmic speech as compared to normal speech.

**Conclusion:** Modulation of connectivity in the cerebellum and prefrontal cortex during rhythmic speech suggests that this fluency-inducing technique activates a compensatory timing system in the cerebellum and potentially modulates top-down motor control and attentional systems. These findings corroborate previous work associating the cerebellum with fluency in adults who stutter and indicate that the cerebellum may be targeted to enhance future therapeutic interventions.

## Introduction

Stuttering is a speech disorder that impacts the production of smooth and timely articulations of planned utterances. Stuttering typically emerges early in childhood and persists over the lifespan for 1% of the population (Craig et al., 2009; Yairi & Ambrose, 1999). Speech of people who stutter (PWS) is characterized by perceptually salient repetitions and prolongations of individual phonemes, as well as abnormal silent pauses at the onset of syllables and words accompanied by tension in the articulatory musculature (Max, 2004). These disfluencies are often accompanied by other secondary behaviors such as eye-blinking and facial grimacing (Guitar, 2014). Along with these more overt characteristics, stuttering also has a severe impact on those who experience it, including increased social anxiety and decreased self-confidence, emotional functioning, and overall mental health (Craig et al., 2009; Craig & Tran, 2006, 2014). Gaining a better understanding of how and why stuttering occurs will help to lead to more targeted therapies and improve quality of life for PWS.

Throughout the years, considerable effort has been made to identify the core pathology underlying stuttering (for reviews, see Max, 2004; Max et al., 2004). More recently, diverse brain imaging modalities have been used to examine how the brains of people who stutter differ from those who do not and how these measures change in different speaking scenarios or following therapy (see Etchell et al., 2018 for a complete literature review). Studies have consistently found that PWS show structural and functional differences in the brain network pertaining to speech initiation and timing (cortico-thalamo-basal ganglia motor loop; Chang & Zhu, 2013; Giraud, 2008; Lu, Peng, et al., 2010) and reduced structural integrity in speech planning areas (left ventral premotor cortex [vPMC] and inferior frontal gyrus [IFG]; Beal et al., 2013, 2015; Chang et al., 2008, 2011; Garnett et al., 2018; Kell et al., 2009; Lu et al., 2012). Functionally, previous work has indicated that during speech, adults who stutter (AWS) have reduced activation in left hemisphere auditory areas (Belyk et al., 2015; Braun et al., 1997; Chang et al., 2009; De Nil et al., 2000, 2008; Fox et al., 1996; Van Borsel et al., 2003) and overactivation in right hemisphere structures (Braun et al., 1997; De Nil et al., 2000; Fox et al., 1996, 2000; Ingham et al., 2000; Van Borsel et al., 2003), which are typically non-dominant for language processing. These studies strongly suggest that stuttering occurs as the result of impaired speech timing, planning, and auditory processing, and that brain structures not normally involved in speech production are potentially recruited to compensate.

In addition to these task activation analyses, previous studies have examined task-based functional connectivity (i.e. activation coupling between multiple brain areas during a speaking task) differences between AWS and ANS. Some studies show reduced connectivity between left IFG and left precentral gyrus in AWS (Chang et al., 2011; Lu et al., 2009), which suggests an impairment in translating speech plans for motor execution (Guenther, 2016). Other studies show group differences in connectivity between auditory, motor, premotor, and subcortical areas ( Chang et al., 2011; Kell et al., 2018; Lu, Chen, et al., 2010; Lu et al., 2009; Lu, Peng, et al., 2010). Results of these task-based connectivity studies, as well as resting-state and structural connectivity studies (e.g., Chang & Zhu, 2013; Sitek et al., 2016), have made it apparent that stuttering behavior is not merely the result of disruptions to one or more separate brain regions, but also differences in the ability for brain regions to communicate with one another during speech.

In addition to examining neural activation in AWS during typical speech, imaging studies have also looked at activation during conditions where AWS speak more fluently. One such condition that has been widely examined behaviorally is the *rhythm effect* in which stuttering disfluencies are dramatically reduced when speakers synchronize their speech movements with rhythmic pacing stimuli (Azrin et al., 1968; Barber, 1940; Hutchinson & Norris, 1977; Stager et al., 1997; Toyomura et al., 2011). These fluency-enhancing effects are robust; they occur regardless of whether the pacing stimulus is presented in the acoustic or visual modalities (Barber, 1940), can be induced even by an imagined rhythm (Barber, 1940; Stager et al., 2003), and occur independently of speaking rate (Davidow, 2014; Hanna & Morris, 1977). Previous studies investigating changes in brain activation during the rhythm effect (Braun et al., 1997; Stager et al., 2003; Toyomura et al., 2011, 2015) have found that during rhythmic speech, both AWS and ANS had increased activation in speech-related auditory and motor regions of cortex as well as parts of the basal ganglia. These activation increases were especially pronounced for AWS as compared to ANS. (Toyomura et al., 2011) also demonstrated that these activation increases occurred in regions displaying under-activation during the normal speaking condition. This suggests that pacing speech along with a metronome improves fluency by “normalizing” under-activation in speech production regions. In light of the functional connectivity studies mentioned previously, characterizing changes in brain connectivity between typical and rhythmically-paced speech could illuminate how external pacing leads to normalized activation in the speech network and, ultimately, fluency.

In the present study, we employed functional MRI during an overt rhythmic sentence-reading task in AWS and ANS to characterize modulation of brain activation and functional connectivity related to the rhythm effect in stuttering. Meta-analyses in neurotypical adults have implicated a common network for rhythmic perceptual and motor timing (Chauvigné et al., 2014; Wiener et al., 2010) involving the cerebellum, basal ganglia, supplementary motor area, and prefrontal cortex, areas which have been integrated into models of rhythmic processing (Teki et al., 2012; Zeid & Bullock, 2019). Therefore, we predict that this network, and its connections with auditory and motor areas normally active during speech production, would be recruited to a larger extent during rhythmic compared to normal speech.

## Method

The current study complied with the principles of research involving human subjects as stipulated by the Boston University institutional review board (protocol 2421E) and the Massachusetts General Hospital human research committee, and participants gave informed consent before taking part. The entire experimental procedure took approximately 2 hours, and subjects received monetary compensation.

### Subjects

Seventeen AWS (12 males/5 females, aged 18-58 years, mean age = 29.8 years, SD = 12.5 years) and seventeen ANS (11 males/6 females, aged 18-49 years, mean age = 28.7 years, SD = 8.1 years) from the greater Boston area were tested. Age was not significantly different between groups (two-sample t-test; *t* = 0.31, *p* = 0.759). Subjects were native speakers of American English who reported normal (or corrected-to-normal) vision and no history of hearing, speech, language, or neurological disorders (aside from persistent developmental stuttering for the AWS). Handedness was measured with the Edinburgh Handedness Inventory (Oldfield, 1971). Using this metric, all AWS were found to be right-handed (scoring greater than 40), but there was more variability among ANS (13 right-handed, 1 left-handed, and 3 ambidextrous). There was a significant difference in handedness score between groups (Wilcoxon rank-sum test; *z* = 2.20, *p* = 0.028); therefore, handedness score was included as a covariate in all group imaging comparisons. For each stuttering participant, stuttering severity was determined using the Stuttering Severity Instrument, Fourth Edition (Riley, 2008); mean score = 23.6, range: 9 to 42; see Table 1 for individual participants). Four additional subjects (3 AWS and 1 ANS) were tested, but they were excluded during data inspection (described below in the *Behavioral Analysis* and *Task Activation fMRI Analysis* sections).

**Table 1:** Demographic and stuttering severity data from adults who stutter. F = female; M = male; SSI-4 = Stuttering Severity Index – Fourth Edition. SSI-Mod = a modified version of the SSI-4 that does not include a subscore related to concomitant movements. Disfluency Rate = the percent of trials containing disfluencies during the Normal speech condition.

| Subject ID | Age | Gender | SSI-4 Composite | SSI-Mod | Disfluency Rate |
| --- | --- | --- | --- | --- | --- |
| AWS01 | 19 | F | 28 | 19 | 0% |
| AWS02 | 22 | F | 31 | 26 | 3.03% |
| AWS03 | 31 | F | 30 | 22 | 3.03% |
| AWS04 | 21 | M | 9 | 7 | 1.92% |
| AWS05 | 58 | M | 14 | 11 | 0% |
| AWS06 | 23 | M | 42 | 29 | 0% |
| AWS07 | 53 | M | 27 | 22 | 0% |
| AWS08 | 44 | M | 20 | 16 | 0% |
| AWS09 | 20 | M | 18 | 15 | 1.52% |
| AWS10 | 22 | M | 27 | 18 | 3.02% |
| AWS11 | 21 | M | 19 | 16 | 6.06% |
| AWS12 | 20 | M | 24 | 14 | 1.52% |
| AWS13 | 29 | M | 33 | 28 | *Missing Data |
| AWS14 | 18 | F | 14 | 11 | 0% |
| AWS15 | 35 | M | 30 | 19 | 0% |
| AWS16 | 42 | M | 22 | 17 | 1.52% |
| AWS17 | 29 | M | 14 | 12 | 0% |

### fMRI Paradigm

Sixteen eight-syllable sentences were selected from the Revised List of Phonetically Balanced Sentences (Harvard Sentences; (*IEEE Recommended Practice for Speech Quality Measurements*, 1969; see Appendix). These sentences, composed of one- and two-syllable words, contain a broad distribution of English speech sounds (e.g. “The juice of lemons makes fine punch”). During a functional brain-imaging session, subjects read aloud the stimulus sentences under two different speaking conditions, one in which individual syllables were rhythmically paced by isochronous auditory beats (i.e., the *rhythm* condition), and one in which syllables were produced using a normal (unmodified) speech rate (*i.e.*, the *normal* condition). For each trial, subjects were presented with eight isochronous tones (1000 Hz, 25ms duration) with a 270 ms interstimulus interval. This resulting rate of approximately 222 beats/min was chosen so that participants’ speech would approximate the rate of the *normal* condition (based on previous estimates of mean speaking rate in English; (Davidow, 2014; Pellegrino et al., 2004). Participants were instructed to refrain from using any part of their body (e.g., finger or foot) to tap to the rhythm.

To avoid confounding interpretation of the BOLD response related to speech production with that of processing the auditory stimulus, the pacing tones were terminated prior to the presentation of the orthographic stimulus. On *rhythm* trials, subjects used the tones to pace their forthcoming speech, while on *normal* trials, they were instructed to disregard the tones and to read the stimuli at a normal speaking rate, rhythm and intonation. During a *rhythm* or *normal* trial, the orthography of a given sentence was presented with the corresponding trial identifier (i.e., “Rhythm” or “Normal”) presented above the sentence. The font color was either blue for *rhythm* and green for *normal* or vice versa, and colors were counterbalanced across subjects. Subjects were instructed to begin reading aloud immediately after the sentence appeared on the screen. In the event that they made a mistake, they were asked to refrain from producing any corrections and remain silent until the next trial. Silent *baseline* trials were also included wherein subjects heard the tones, and saw a random series of typographical symbols (e.g. ‘+\^ &$/[|\ $=[ [)*% /-@ \| -%-/’) clustered into word-like groupings (matched to stimulus sentences); subjects refrained from speaking during these trials.

Subjects participated in a behavioral experiment (not reported here) prior to the imaging experiment that gave them experience with the speech stimuli and the task. The time between this prior exposure and the present experiment ranged from 0 to 424 days. Immediately prior to the imaging session, subjects practiced each sentence under both conditions until they demonstrated competence with the task and sentence production. Subjects also completed a set of six practice trials in the scanner prior to fMRI data collection. To control basic speech parameters across conditions and groups, subjects were provided with performance feedback on their overall speech rate and loudness during practice only. Following this practice set, subjects completed between two and four experimental runs of test trials depending on time constraints (29 completed four, 4 completed three, 1 completed two). During the experimental session, verbal feedback was provided between runs if subjects consistently performed outside of the specified speech rate (220 ms to 320 ms mean syllable duration). Each run consisted of 16 *rhythm* trials, 16 *normal* trials, and 16 *baseline* trials, pseudo-randomly interleaved within each run for each subject. All trials were audio-recorded for later processing.

### Data Acquisition

MRI data for this study were collected at two locations: the Athinoula A. Martinos Center for Biomedical Imaging at the Massachusetts General Hospital (MGH), Charlestown Campus (9 AWS, 9 ANS) and the Cognitive Neuroimaging Center at Boston University (BU; 8 AWS, 8 ANS). At MGH, images were acquired with a 3T Siemens Skyra scanner and a 32-channel head coil, while a 3T Siemens Prisma Scanner with a 64-channel head coil was used at BU. At each location, subjects lay supine in the scanner and functional volumes were collected using a gradient echo, echo planar imaging BOLD sequence (repetition time [TR] = 11.5 s, acquisition time = 2.47 s, TE = 30 ms, Flip Angle = 90°). Each functional volume covered the entire brain and was composed of 46 axial slices (64 × 64 matrix) acquired in interleaved order and accelerated using a simultaneous multislice factor of 3 with a 192 mm field of view. The in-plane resolution was 3.0 × 3.0 mm^2^, and slice thickness was 3.0 mm with no gap. Two “dummy” scans were included at the beginning of each run to ensure equilibrium in the magnetic field prior to data collection. Additionally, a high-resolution T1-weighted whole-brain structural image was collected from each participant to anatomically localize the functional data (MPRAGE sequence, 256 × 256 × 176 mm^3^ volume with a 1 mm isotropic resolution, TR = 2.53 s, inversion time = 1100 ms, echo time = 1.69 ms, flip angle = 7°).

Functional data were acquired using a sparse image acquisition paradigm (Eden et al., 1999; Hall et al., 1999) that allowed participants to produce the target sentences during silent intervals between volume acquisitions. Volumes were acquired 5.7-8.17 s after stimulus presentation to ensure a 4-6 second delay between the middle of sentence production and the acquisition, in alignment with the delay in the peak of the task-related blood oxygen-level-dependent (BOLD) response (Belin et al., 1999). By scanning after speech production has ended, this paradigm reduces head motion-induced scan artifacts, eliminates the influence of scanner noise on speaker performance, and allows subjects to perceive their own self-generated auditory feedback in the absence of scanner noise (e.g., Gracco et al., 2005). A schematic representation of the trial structure and timeline is shown in Figure 1.

**Figure 1:**
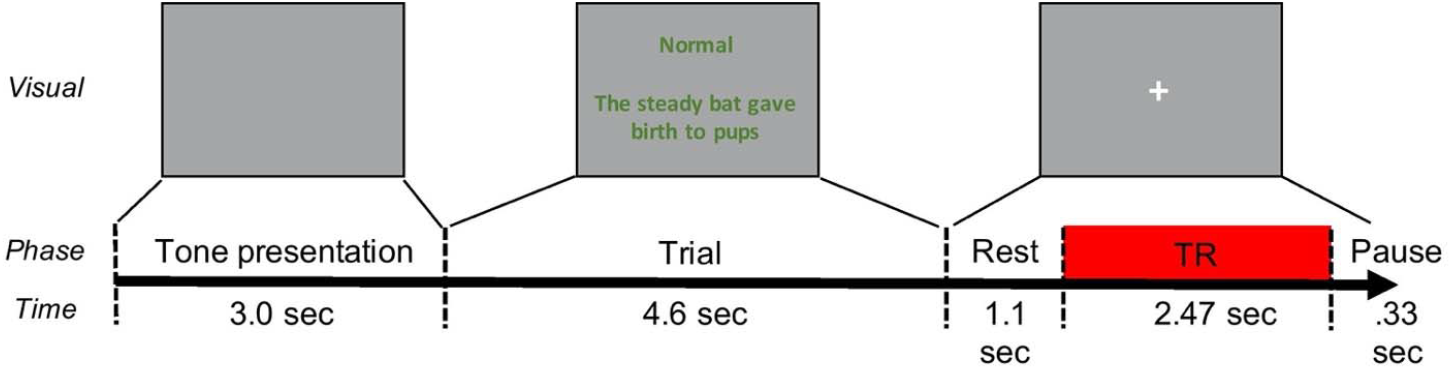
Schematic diagram illustrating the temporal structure of stimulus presentation during functional data acquisition. At the start of each trial, isochronous tone sequences were presented for 3.0 seconds. The visual stimulus then appeared and remained on screen for 4.6 seconds. 1.1 seconds after stimulus offset, a whole-brain volume was acquired. The next trial started 0.33 seconds after data acquisition was complete. TR = repetition time.

Visual stimuli were projected onto a screen viewed from within the scanner via a mirror attached to the head coil. Auditory stimuli were delivered to both ears through Sensimetrics model S-14 MRI-compatible earphones using Matlab (The MathWorks, Natick, MA). Subjects’ utterances were transduced with a Fibersound model FOM1-MR-30m fiber-optic microphone, sent to a laptop (Lenovo ThinkPad W540), and recorded using Matlab. Subjects took a short break after completing each run.

### Behavioral Analysis

An automatic speech recognition engine was used to objectively measure how accurately subjects aligned their syllables to the metronome beats. Specifically, the open-source large-vocabulary continuous speech recognition engine *Julius* (Lee & Kawahara, 2009) was used in conjunction with the free *VoxForge* American English acoustic models (voxforge.org) to perform phoneme-level alignment on the sentence recordings. This resulted in phoneme boundary timing information for every trial. A researcher manually inspected each trial to ensure correct automatic detection of phoneme boundaries. Any trials in which the subject made a reading error, a condition error (i.e. spoke rhythmically when they were cued to speak normally or vice versa), or a disfluency categorized as a stutter by a licensed speech-language pathologist were eliminated from further behavioral analysis. One ANS that made consistent condition errors was eliminated from further analysis. One AWS was eliminated from further analysis due to an insufficient number of fluent trials during the *normal speech* condition (6/64 attempted). Additionally, following the error trial elimination step, behavioral data from AWS13 were deleted due to a technical error, so only 16 AWS are included in the behavioral analyses.

To evaluate whether there was a fluency-enhancing effect of rhythmic pacing, the percentage of trials eliminated due to stuttering in the AWS group was compared between the two speaking conditions using a non-parametric Wilcoxon signed-rank test. Measures of the total sentence duration and intervocalic timing from each trial were also extracted to determine the rate and isochronicity of each production. Within a sentence, the average time between the centers of the eight successive vowels was calculated to determine the intervocalic interval (IVI). The reciprocal (1/IVI) was then calculated, resulting in a measure of speaking rate in units of IVIs per second. The coefficient of variation for intervocalic intervals (CV-IVIs) was also calculated by dividing the standard deviation of IVIs divided by the mean IVI. A higher CV-IVI indicates higher variability of IVI, while a CV-IVI of 0 reflects perfect isochronicity. Rate and CV-IVI were compared between groups and conditions using a mixed design ANOVA. A Bonferroni correction was applied across these two analyses to account for testing these related measures.

### Task Activation fMRI Analysis

#### Preprocessing

Following data collection, all images were processed through two preprocessing pipelines: a surface-based pipeline for cortical activation analyses and a volume-based pipeline for subcortical and cerebellar analyses. For the surface-based pipeline, functional images from each subject were simultaneously realigned to the mean subject image and unwarped (motion-by-inhomogeneity interactions) using SPM12’s realign and unwarp procedure (Andersson et al., 2001). Outlier scans were detected with Artifact Detection Tools (ART; https://www.nitrc.org/projects/artifact_detect/) based on motion displacement (scan-to-scan motion threshold of 0.9 mm) and mean signal change (scan-to-scan signal change threshold of 5 standard deviations above the mean). Functional images from each subject were then coregistered with their high-resolution T1 structural images and resliced using SPM12’s inter-modal registration procedure with a normalized mutual information objective function. The structural images were segmented into white matter, grey matter, and cerebrospinal fluid, and cortical surfaces were reconstructed using the FreeSurfer image analysis suite (freesurfer.net; Fischl et al., 1999). Functional data were then resampled at the location of the FreeSurfer fsaverage tessellation of each subject-specific cortical surface.

For the volume-based pipeline, functional volumes were realigned and unwarped, centered, and run through ART as described for the surface-based pipeline. Functional volumes were then simultaneously segmented and normalized directly to Montreal Neurological Institute (MNI) space using SPM12’s combined normalization and segmentation procedure (Ashburner & Friston, 2005). A mask was then applied such that only voxels within the brain were submitted to subsequent analyses. The original T1 structural image from each subject was also centered, segmented and normalized using SPM12.

Following preprocessing, two AWS were eliminated from subsequent analyses; one due to excessive head motion in the scanner (>1.5mm average scan-to-scan motion) and one due to structural brain abnormalities.

#### First-level Analysis

After preprocessing, BOLD responses were estimated for each subject using a general linear model (GLM) in SPM12. Because images were collected in a sparse sequence with a relatively long TR, the BOLD response for each trial (event) was modeled as an individual epoch. The model included regressors for each of the conditions of interest: *normal speech, rhythm speech*, and *baseline.* Trials that contained reading errors, condition errors, or disfluencies were modeled as a single separate condition of non-interest. Condition regressors were collapsed across runs to maximize power while controlling for potential differences in the number of trials produced without errors or disfluencies. For each run, regressors were added to remove linear effects of time (e.g. signal drift, adaptation) in addition to six motion covariates (taken from the realignment step) and a constant term. Additional regressors were added to remove the effects of acquisitions with excessive scan-to-scan motion or global signal change (taken from the artifact detection step, described above). The first-level model regressor coefficients for the three conditions of interest were estimated at each surface vertex and subcortical voxel, then averaged within anatomical regions of interest (ROIs; see below). The mean *normal speech* and *rhythm speech* coefficients were then contrasted with the *baseline* condition within each ROI to yield contrast effect-size values for the two contrasts of interest (*Normal – Baseline* and *Rhythm – Baseline)* in all ROIs.

#### Region-of-Interest Definition

Cortical ROIs were labeled according to a modified version of the SpeechLabel atlas previously described in (Cai et al., 2014); the atlas divides the cortex into macro-anatomically defined ROIs specifically tailored for studies of speech. Labels are applied by mapping the atlas from the FreeSurfer *fsaverage* cortical surface template to each individual surface reconstruction.

Subcortical and cerebellar ROIs were extracted from multiple atlases. Thalamic ROIs were extracted from the mean atlas of thalamic nuclei described by (Krauth et al., 2010). Basal ganglia ROIs were derived from the non-linear normalized probabilistic atlas of basal ganglia (ATAG) described by (Keuken et al., 2014). Each was ROI was thresholded at a minimum probability threshold of 33% and combined in a single labeled volume in the atlas’s native space (the MNI104 template). Cerebellar ROIs were derived from the SUIT 25% maximum probability atlas of cerebellar regions (Diedrichsen, 2006; Diedrichsen et al., 2009, 2011). Each atlas was non-linearly registered to the SPM12 MNI152 template and then combined into a single labeled volume.

#### Second-Level Group Analyses

Two sets of analyses were carried out to detect activation differences across groups and conditions: hypothesis-based primary analyses, and exploratory secondary analyses. The primary second-level analyses were carried out on a small set of hypothesis-based *a priori* ROIs (see Figure 2). These included regions belonging to the cortico-basal ganglia-thalamo-cortical motor loop (Guenther, 2016), meta-analyses of rhythmic perceptual and motor timing (Chauvigné et al., 2014; Wiener et al., 2010), and prior neuroimaging studies examining the rhythm effect in stuttering (Stager et al., 2003; Toyomura et al., 2011). Statistical corrections were applied for the number of ROIs tested. The following cortical ROIs in the SpeechLabel atlas were grouped to test our hypotheses: ventral and mid primary motor cortex (MC), ventral and mid premotor cortex (PMC), supplementary motor area and pre-supplementary motor area (SMA), posterior superior temporal gyrus and planum temporale (pSTg), and ventral and dorsal inferior frontal gyrus pars opercularis (IFo). By grouping the ROIs, we better match the extent of areas shown to be involved in rhythm processing/stuttering in prior reports and increase the sensitivity of our analyses by reducing the number of ROIs.

**Figure 2:**
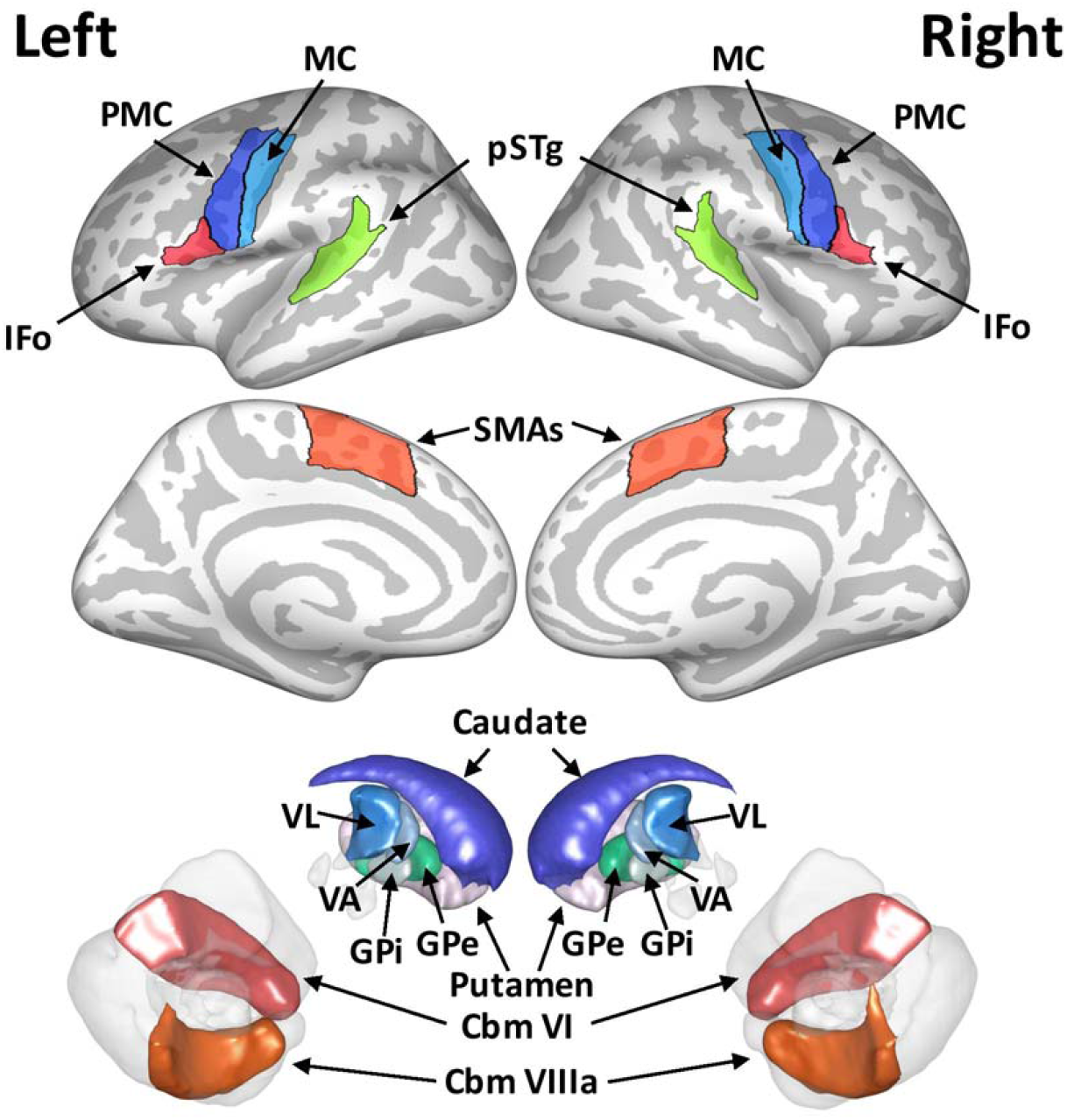
Regions of interest included in the primary hypothesis-based analysis. Cortical regions are displayed on an inflated cortical surface, while subcortical and cerebellar regions are rendered in 3-D volume space. IFo = grouped dorsal and ventral inferior frontal gyrus pars operularis, PMC = grouped ventral and mid premotor cortex, MC = grouped ventral and mid motor cortex, pSTg = grouped posterior superior temporal gyrus, SMAs = grouped supplementary motor areas, VL = ventrolateral thalamic nucleus, VA = ventroanterior thalamic nucleus, GPi = internal portion of globus pallidus, GPe = external portion of globus pallidus, Cbm VI = cerebellum lobule VI, Cbm VIIIa = cerebellum lobule VIIIa.

Additional exploratory analyses were performed to determine if activation from other brain regions active during speech production was also modulated by group or condition. For each exploratory analysis, results are reported if they have a *p*-value less than 0.05, uncorrected. To determine this set of regions, second-level random effects analyses were performed on first-level contrast effect sizes in all ROIs for each group separately. Regions with significant positive activation (thresholded at one-*sided p* < 0.05, and corrected for multiple comparisons using a false discovery rate correction [FDR; Benjamini & Hochberg, 1995] within each contrast) in any of these four contrasts were included in subsequent analyses (see Supplementary Figure 2 and Supplementary Figure 3 for the complete list).

Group activation differences were examined in the two speech conditions compared to baseline (*Normal – Baseline, Rhythm – Baseline)* as well as the *Group × Condition Interaction.* Additionally, differences between the two speech conditions (*Rhythm – Normal*) were examined in each group separately. These group and condition effects were determined using a GLM. Average subject motion was added as a regressor of non-interest for all analyses. In addition, to account for differences across the two data collection sites, an additional regressor of non-interest was included for all analysis. Due to significant difference in handedness between the two groups (see Subjects section above), handedness score was also included as a regressor of non-interest for between-group and interaction analyses. Finally, to control for stuttering severity, a modification of the SSI-4 score, heretofore termed “SSI-Mod,” was included as another regressor of non-interest in the between-group and interaction analyses. SSI-Mod removes the secondary concomitants subscore from each subject’s SSI-4 score, thus focusing the measure on speech-related function. The SSI-Mod and SSI-4 composite scores for each subject are included in Table 1. Additional regression analyses were carried out to determine whether stuttering severity, measured by the SSI-Mod, or disfluencies occurring during the experiment were correlated with task activation. Because very few disfluencies occurred during the rhythm condition, we were only able to calculate the correlation between the percentage of disfluencies occurring during *normal* trials (“Disfluency Rate”) and the *Normal - Baseline* activation. Note that because trials containing disfluencies were regressed out of the first-level effects, correlations with Disfluency Rate are capturing activation related to the *propensity* to stutter and not disfluent speech itself. The primary analyses were performed using a strict statistical correction of *p_FDR_* < 0.05, while the exploratory analyses were performed using an uncorrected alpha level of 0.05.

### Functional Connectivity Analysis

#### Preprocessing and analysis

Seed-based functional connectivity analyses (SBC) were carried out using the CONN toolbox (Whitfield-Gabrieli & Nieto-Castanon, 2012). The same preprocessed data used for the task activation analysis were used for the functional connectivity analysis. The seeds for this analysis comprised the same “speech production” ROIs used in the exploratory task activation analysis, defined either in *fsaverage* surface (cortical) or MNI volume (subcortical) space. The BOLD time series was averaged within seed ROIs. To include connections between the speech production network and other regions that potentially have a moderating effect on this network, the target area in this analysis was extended to the whole brain. The target functional volume data were smoothed using an 8 mm full-width half maximum Gaussian smoothing kernel. Following preprocessing, an aCompCor (Behzadi et al., 2007) denoising procedure was used to eliminate extraneous motion, physiological, and artifactual effects from the BOLD signal in each subject. In each seed ROI and every voxel in the smoothed brain volume, denoising was carried out using a linear regression model (Nieto-Castañón, 2020) that included 5 white matter regressors, 5 CSF regressors, 6 subject-motion parameters plus their first-order temporal derivatives, scrubbing regressors to remove the effects of outlier scans (from artifact detection, described above), as well as separate regressors for each run/session (constant effects and first-order linear-trends), task condition (main and first-order derivative terms), and error trials. No band-pass filter was applied in order to preserve high-frequency fluctuations in the residual data.

For each participant, a generalized PsychoPhysiological Interaction (gPPI; McLaren et al., 2012) analysis was implemented using a multiple regression model, predicting the signal in each target voxel with three sets of regressors: a) the BOLD time series in a seed ROI, characterizing baseline connectivity between a seed ROI and each target voxel; b) the main effects of each of the task conditions (*normal, rhythm*, and *baseline*), characterizing direct functional responses to each task in the target voxel; and c) their seed-time-series-by-task interactions (PPI terms) characterizing the relative changes in functional connectivity strength associated with each task. Second-level random effects analyses were then used to compare these interaction terms within and between groups and conditions, specifically the *Rhythm - Normal* contrast in AWS and ANS and the *Group × Condition* interaction. The same regressors of non-interest used in the task activation analyses were included here as well. For each comparison, separate analyses were run from the 103 seed ROIs to the whole brain. Within each analysis, a two-step thresholding procedure was used; voxels were thresholded at *p* < 0.001, followed by a cluster-size threshold of *p_FDR_* < 0.05. To control for family-wise error across the 103 separate seed-to-voxel analyses, a within-comparison Bonferroni correction was applied so that only significant clusters with *p_FDR_* < 0.000485 (0.05/103) survived the threshold.

### Results

#### Behavioral Analysis

Stuttering occurred infrequently over the course of the experiment, with 7 out of 16 AWS producing no disfluencies. There was, however, a significantly lower percentage of disfluent trials in the *rhythm* condition (0.38%) compared to the *normal* condition (1.35%; *W* = 42, *p* = 0.023; see Figure 3). There was no group *×* condition interaction or group main effect on speaking rate but there was a significant main effect of condition with *normal speech* (3.977 IVI/sec) produced at a faster rate than *rhythmic speech* (3.460 IVI/sec; *F*(1,31) = 37.8, *p_FWE_* < 0.001). For isochronicity, there was no main effect of group or group × condition interaction. There was a significant main effect of condition, where subjects had a lower CV-IVI (greater isochronicity) in the *rhythm* condition (0.25) than the *normal* condition (0.13; *F*(1,31) = 503.3, *p_FWE_* < 0.001).

**Figure 3:**
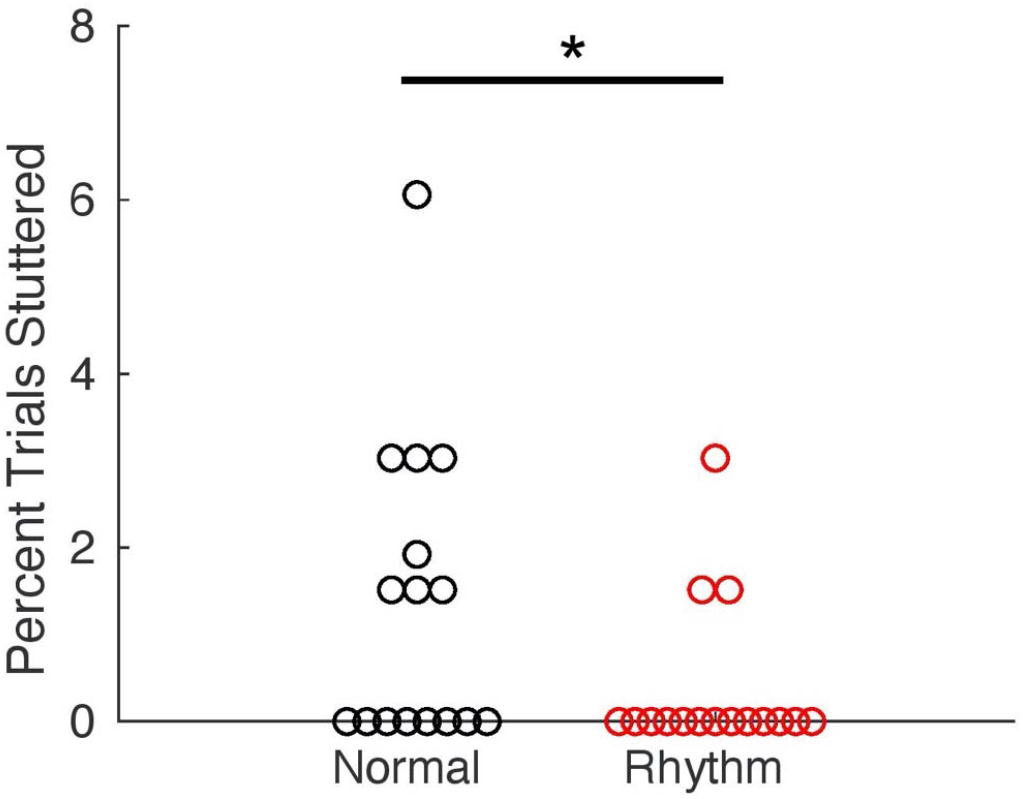
Comparison of dysfluencies between the normal and rhythm conditions for AWS. Circles represent individual participants. *p < 0.05.

#### Task Activation fMRI Analysis

The cortical results of the *Normal* - *Baseline* and *Rhythm - Baseline* contrasts in each group are presented in Supplementary Figure 1. The set of 103 cortical and subcortical ROIs that were significant in at least one of those contrasts and used for subsequent exploratory analyses is illustrated in Supplementary Figures 2 and 3.

For the primary analysis, ANS had greater activation in the *Rhythm* condition compared to the *Normal* condition in left grouped supplementary motor areas (SMAs), posterior superior temporal gyrus (pSTG), ventro-anterior thalamus (VA), and ventro-lateral thalamus (VL), and in right grouped premotor cortex (PMC), caudate nucleus (Caud), and VA (*p_FDR_* < 0.05, see Table 2 and Figure 4). No significant differences were found between conditions in AWS. For the complete exploratory results, see Supplementary Table 1 and Supplementary Figure 4. Notably, eight exploratory ROIs survived an FDR statistical correction: left planum temporale (PT), pre-supplementary motor area (preSMA), superior parietal lobule (SPL), anterior insula (aINS), planum polare (PP), supplementary motor area (SMA), VA, and right ventral premotor cortex (vPMC). For the same exploratory contrast, AWS showed increased activity in left PT, and right ventral inferior frontal gyrus pars opercularis (vIFo) in the *Rhythm* condition, and decreased activation in right anterior dorsal superior temporal sulcus (adSTs) and cerebellar vermis lobule VIIIb. To explore whether the failure of these effects to survive the corrected significance threshold was due to overall greater variability among AWS participants, we averaged *Rhythm - Normal* effects across all exploratory ROIs and performed Levene’s test for equality of variances. AWS had significantly larger variance across subjects (*F* = 3.42, *p* = 0.019).

**Table 2:** Primary regions-of-interest with activation differences between the rhythm and normal conditions for ANS and AWS (p < 0.05). * indicates regions that survive a significance threshold of pFDR < 0.05 for their respective analyses, unc = uncorrested, SMA = grouped supplementary motor areas, PMC = grouped ventral and mid premotor cortex, pSTg = grouped posterior superior temporal gyrus and planum temporale, VA = ventraoanterior thalamic nucleus, VL = ventral lateral thalamic nucleus.

| ROI | Hemisphere | *t-value* | *p-unc* |
| --- | --- | --- | --- |
| *ANS, Rhythm > Normal* |  |  |  |
| SMA<sub>s</sub> | Left | 4.68 | 0.0004* |
| PMC | Right | 3.22 | 0.0062* |
| pSTg | Left | 2.94 | 0.0108* |
| VA | Left | 3.88 | 0.0017* |
|  | Right | 2.98 | 0.0098* |
| VL | Left | 3.59 | 0.0030* |
| Caudate | Right | 3.72 | 0.0023* |

**Figure 4:**
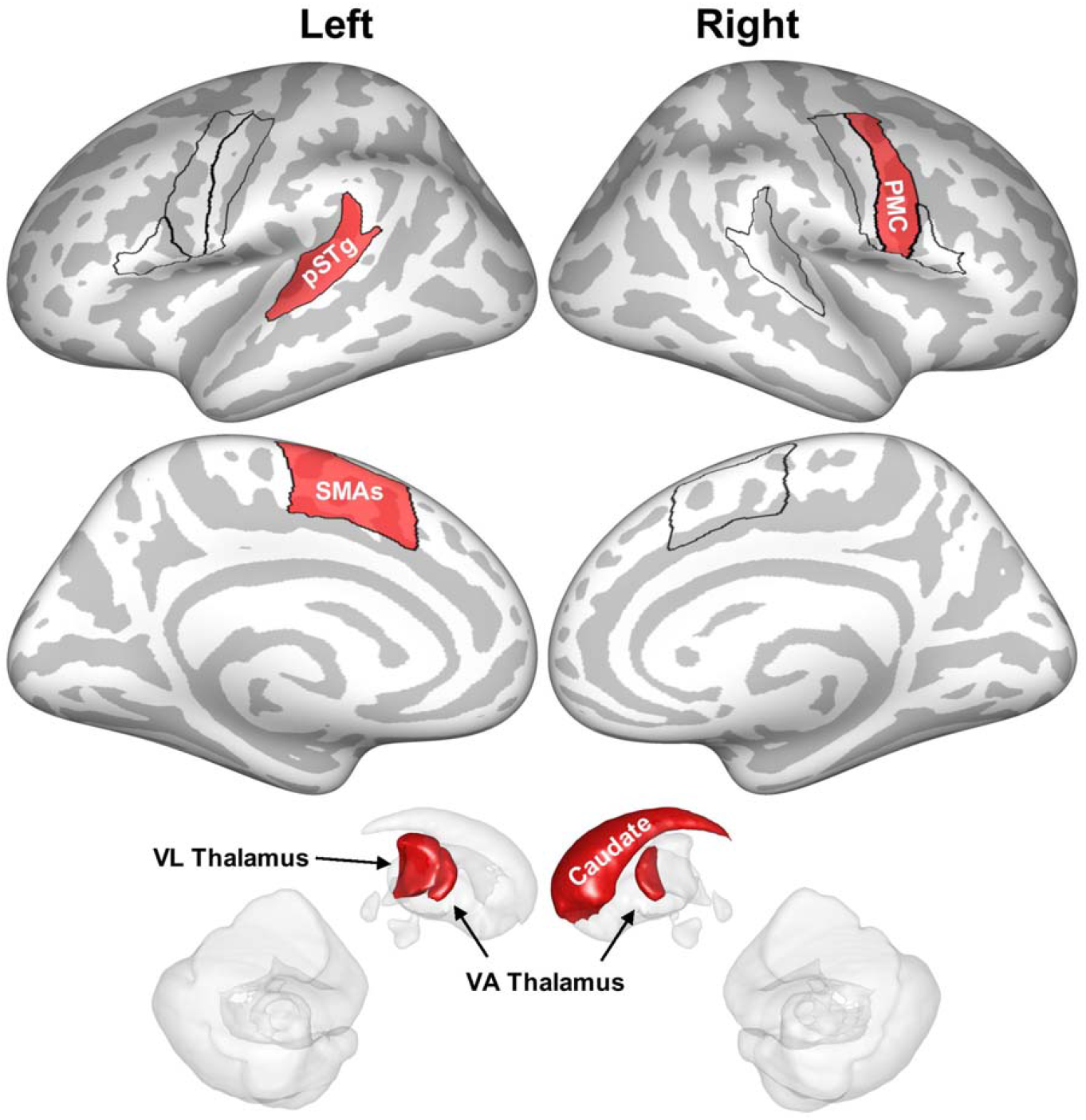
Primary regions-of-interest (ROIs) significantly more active during the rhythmic condition than the normal condition for ANS in the primary analysis (pFDR < 0.05) are highlighted in red and plotted on an inflated cortical surface or on a 3-D rendering of subcortical structures. Black outlines indicate cortical ROIs included in the primary analysis (as in Figure 2). pSTg = posterior superior temporal gyrus, SMAs = grouped supplementary motor areas, PMC = grouped ventral and mid premotor cortex, pSTg = grouped posterior superior temporal gyrus and planum temporale, VA = ventro-anterior, VL = ventro-lateral.

For the primary analysis, no significant differences were found between groups for either *Normal* - *Baseline* or *Rhythm* - *Baseline.* In the exploratory analysis, AWS had decreased activation in left anterior frontal operculum (aFO; *p* = 0.009) and the internal portion of the globus pallidus (GPi; *p* = 0.047), as well as midline cerebellar vermis VIIIb (*p* = 0.038), in the *Rhythm - Baseline* contrast compared to ANS.

In our primary analysis, no ROIs showed a significant interaction between groups and conditions. In the follow-up exploratory analysis, an interaction was found in five ROIs (see Supplementary Table 2 and Supplementary Figure 5): left PP, aFO, cerebellar lobule VIIIa (Cbm VIIIa), and the external portion of the globus pallidus (GPe), and midline cerebellar vermis VIIIb (*p* < 0.05). In all cases, ANS had increased activation in the *Rhythm* condition compared to *Normal*, while AWS showed no change or a decrease.

#### Brain-Behavior Correlation Analyses

In our primary analysis, no significant correlation was found between SSI-Mod and *Normal - Baseline* or *Rhythm - Baseline* in any ROI when correcting for multiple comparisons. There were, however, significant positive correlations between Disfluency Rate and *Normal – Baseline* activation in left VA and VL as well as right VL (Table 3).

**Table 3:** Primary regions-of-interest with significant correlations between severity measures and speech activation in AWS (p < 0.05). * indicates regions that survive a significance threshold of pFDR < 0.05 for their respective analyses, unc = uncorrested, VA = ventroanterior thalamic nucleus, VL = ventrolateral thalamic nucleus.

| ROI | Hemisphere | *t-value* | *p-unc* |
| --- | --- | --- | --- |
| *Normal-Baseline Correlation with Disfluency Rate* |  |  |  |
| VA | Left | 4.15 | 0.0013* |
| VL | Left | 3.43 | 0.0049* |
|  | Right | 3.44 | 0.0049* |

Exploratory results can be found in Supplementary Table 3. Of note, positive correlations were found between SSI-Mod and activation in bilateral premotor and frontal opercular cortex and negative correlations were found in left anterior auditory cortex. In addition, positive correlations between Disfluency Rate and *Normal - Baseline* were found in right parasylvian regions and bilateral putamen.

#### Functional Connectivity Analyses

##### Within group

Within the AWS group, seven connections were significantly stronger in the *Rhythm* condition as compared to the *Normal* condition (*p_FDR_* < 0.000485), all involving the cerebellum (see Table 4 and Figure 5). Both left and right cerebellar lobule VIIIa displayed greater connectivity with clusters in bilateral orbitofrontal cortex (OFC; two distinct clusters with right cerebellar lobule VIIIa [clusters 3 and 4 in Figure 5], and one cluster with left cerebellar lobule VIIIa straddling the midline [cluster 2]), and right cerebellar lobule VIIb had greater connectivity in an overlapping region of right OFC (cluster 1). Right dentate nucleus showed an increase in connectivity with one cluster covering medial cerebellar lobule VI and Crus I (cluster 5), and a second cluster in right lateral cerebellar lobule VI and Crus I (cluster 6). Finally, there was increased connectivity between cerebellar vermis Crus II and a cluster in the superior cerebellum more anteriorly (cluster 7). In all cases, there was either a negative relationship or no relationship during the *Normal* condition, and a positive relationship during the *Rhythm* condition. To determine whether these differences were specific to AWS, a *post hoc* analysis found that these connections did not reach significance in the ANS group, even using an uncorrected alpha level of 0.05. Instead, ANS had different connections that were significantly stronger during *Rhythm* speech compared to *Normal:* between left VA and a cluster in right occipital cortex (OC) and fusiform gyrus (FG); and right preSMA and a cluster at the junction of left SPL, precuneus (PCN), and OC. There was also a decrease in connectivity between left substantia nigra (SN) and a cluster in left OC (see Supplementary Figure 6).

**Table 4:**
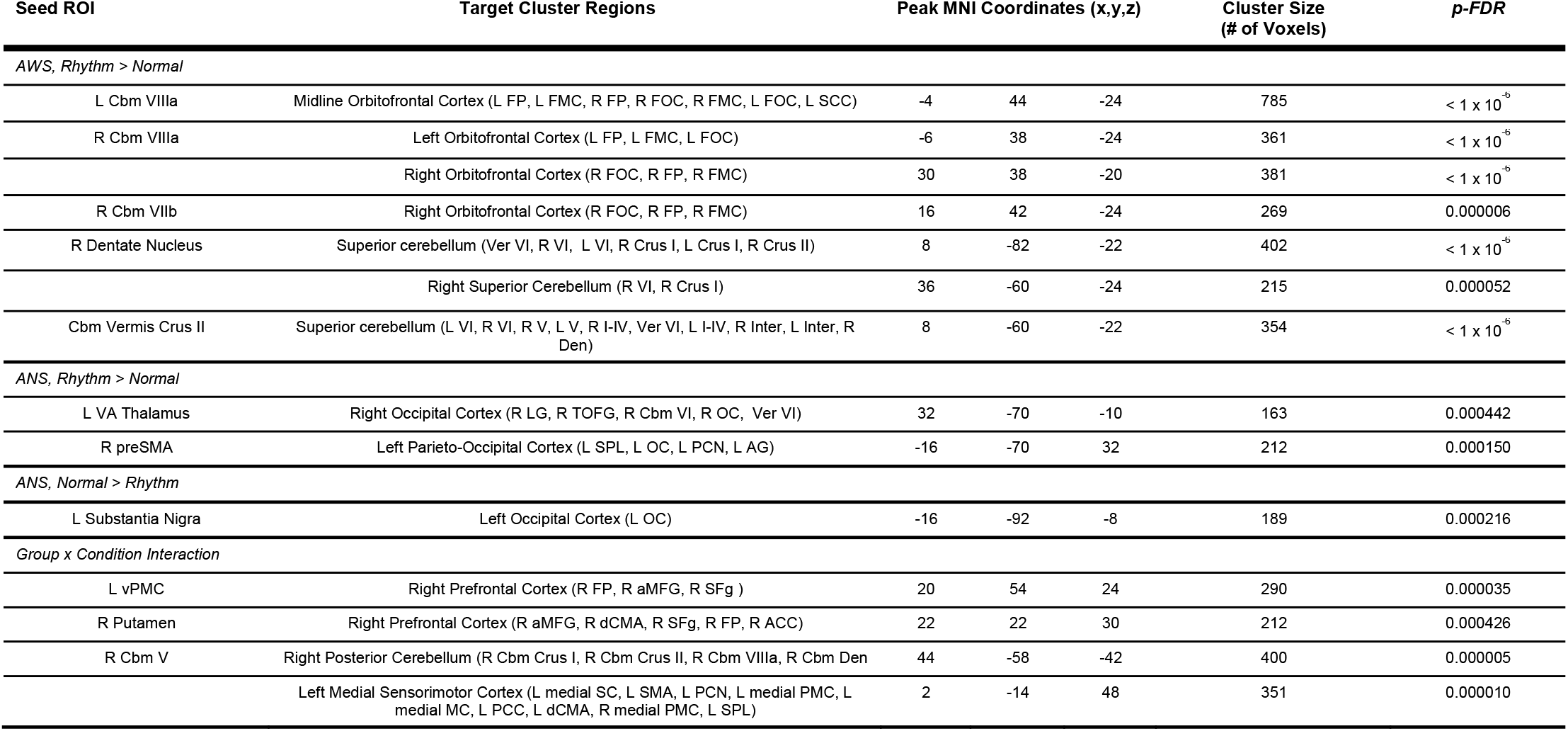
Functional connectivity analysis results. ROI = region-of-interest, MNI = Montreal Neurological Institute, FDR = false discovery rate, L = left, R = right, Cbm = cerebellum, FP = frontal pole, FMC = fronto-medial cortex; FOC = fronto-orbital cortex, SCC = subcallosal cortex, Inter = interposed nucleus, Den = dentate nucleus, vCMA = ventral cingulate motor area, dCMA = dorsal cingulate motor area, ACC = anterior cingulate cortex, OC = occipital cortex, LG = lingual gyrus, TOFG = temporo-occipital fusiform gyrus, SPL = superior parietal lobule, PCN = precuneus AG = angular gyrus, aMFG = anterior middle frontal gyrus, SFG = superior frontal gyrus, preSMA = presupplementary motor area, MC = primary motor cortex, SC = somatosensory cortex, SMA = supplementary motor area, PCC = posterior cingulate cortex, SPL = superior parietal lobule.

**Figure 5:**
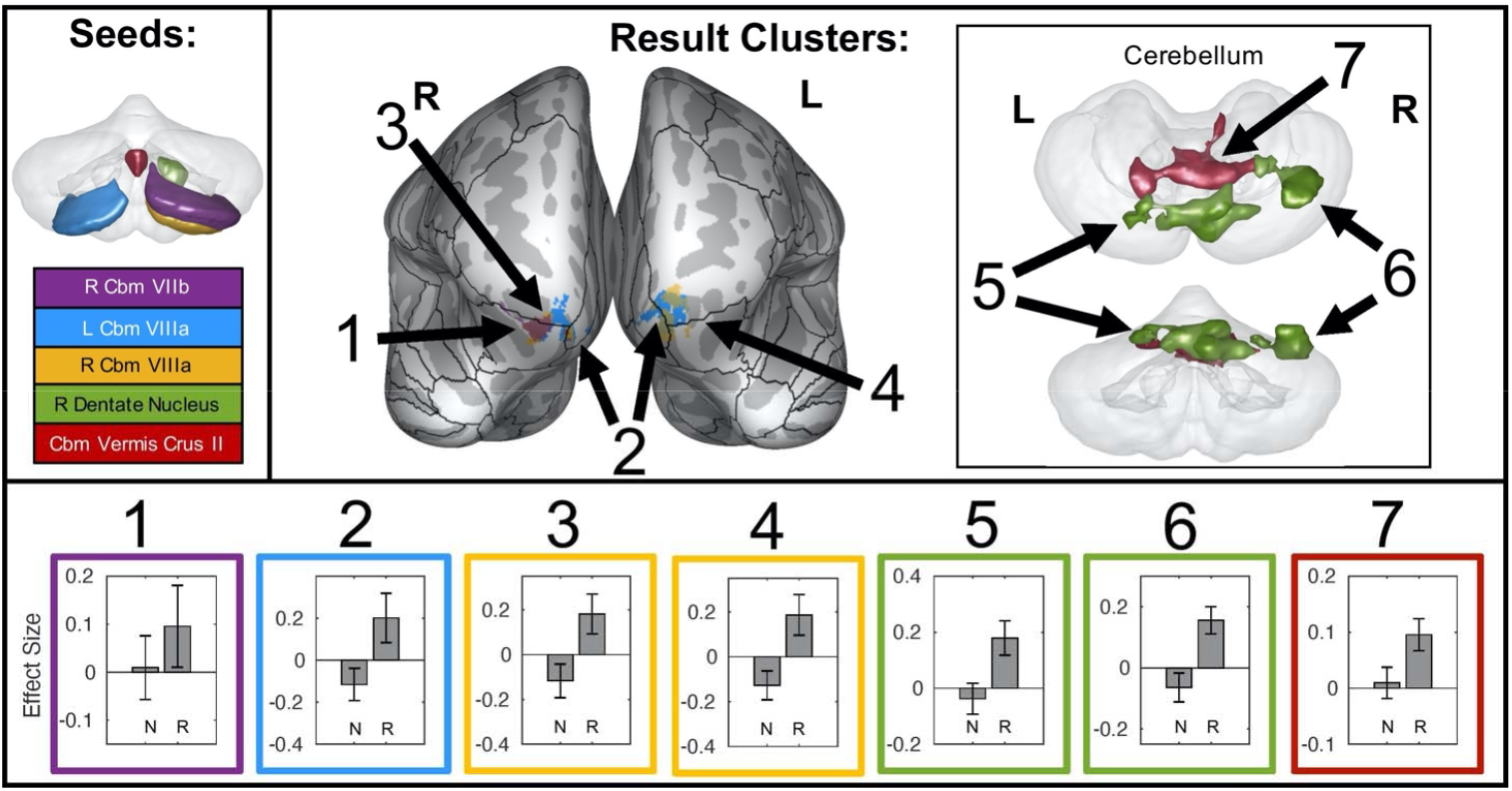
A summary of functional connections that are significantly different between the normal and rhythm conditions in AWS. Seed regions for these connections are indicated in the upper left corner on a transparent 3D rendering of the cerebellum (viewed posteriorly), and colors in the rest of the figure refer back to these seed regions. Seven target clusters (representing 7 distinct connections) are displayed in the upper right portion of the figure. Target clusters 1-4 are projected onto an inflated surface of cerebral cortex, along with the full cortical ROI parcellation of the SpeechLabel atlas described in Cai et al. (2014). Target clusters 5, 6 and 7 are displayed on a transparent 3D rendering of the cerebellum (top view: superior; bottom view: posterior). The bottom portion of the figure shows the connectivity effect sizes in the normal and rhythm conditions for each connection. Error bars indicate 90% confidence intervals. N = normal, R = rhythm, L = left, R = right, Cbm = cerebellum.

##### Group × Condition Interaction

There were four connections that showed a significant interaction between group and speech condition (*Normal* and *Rhythm;* see Figure 6). Connections that were lower in the *Rhythm* condition for AWS and greater in this condition for ANS included: right cerebellar lobule V to left medial rolandic cortex and posterior SMA (result cluster labeled 1 in bottom-left panel of Figure 6); left putamen to right aMFG (extending to right medial cortex; cluster 2); and left vPMC to left frontal pole (FP) and anterior middle frontal gyrus (aMFG; cluster 3). A connection that was greater in the *Rhythm* condition for AWS and lesser in this condition for ANS was between right cerebellar lobule V to right cerebellar lobule Crus I, Crus II, and dentate nucleus (cluster 4). Simple effects from each group and condition are shown in the bottom panel of Figure 6. Based on the results that showed increased connectivity for AWS between different parts of the cerebellum during rhythmic speech, we performed a test comparing average pairwise connectivity among all 21 cerebellar ROIs active during speech. This test revealed that these ROIs show a significant group × condition interaction (*t* = 2.90*, p* = 0.004), driven by an increase in connectivity for AWS from *Normal* to *Rhythm (t* = 3.94,*p* < 0.001) and a non-significant decrease in connectivity for ANS (*t* = −1.23,*p* = 0.880).

**Figure 6:**
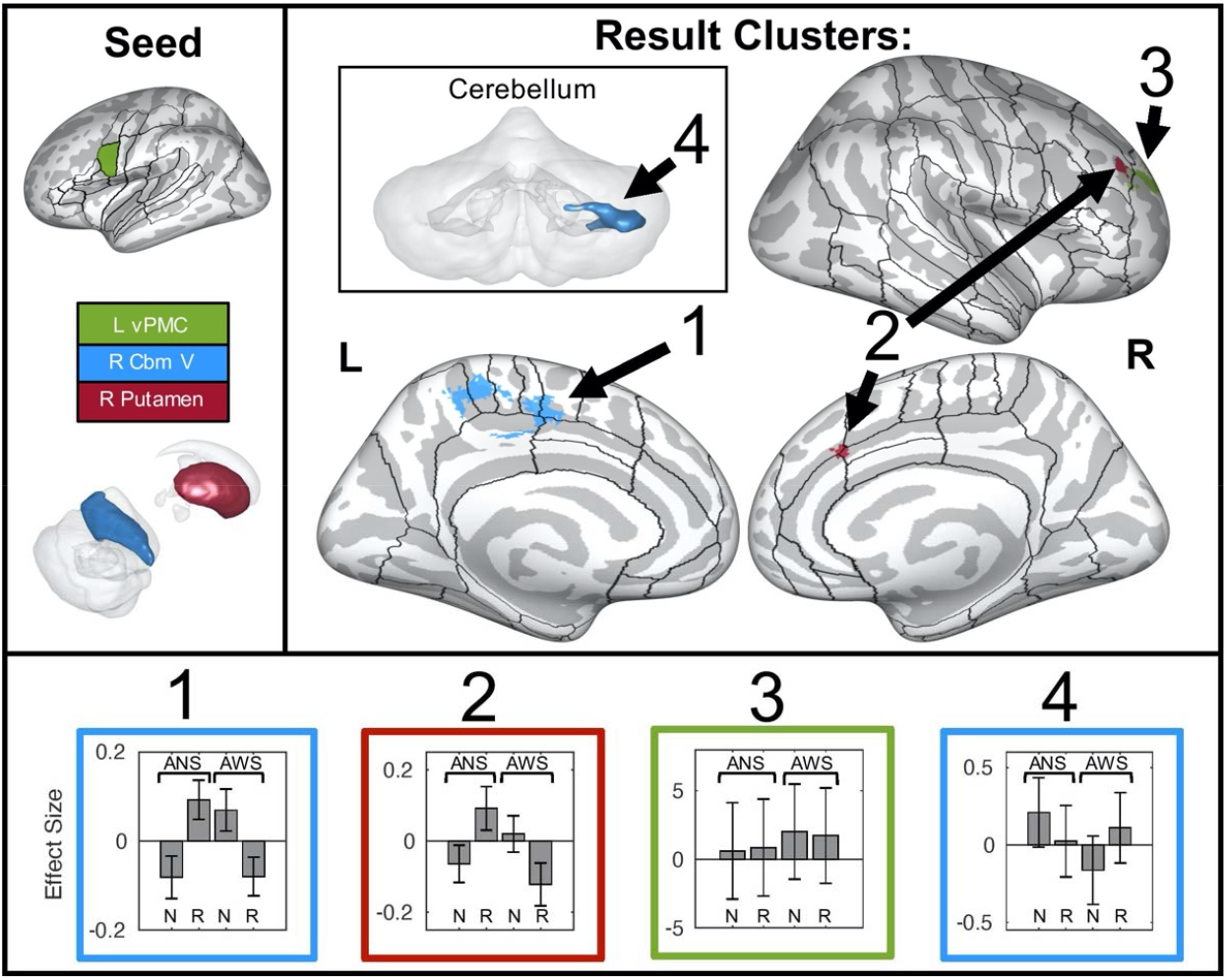
A summary of functional connections that show significant interactions between group and condition. Seed regions for these connections are indicated in the upper left panel on an inflated cortical surface (top; ROIs are as in Figure 2) or on a transparent 3D rendering of the cerebellum and subcortical structures viewed from the right (bottom). Colors in the rest of the figure refer back to these seed regions. Four target clusters (representing 4 distinct connections) are displayed in the upper right portion of the figure. Target clusters 1, 2, and 3 are projected onto an inflated surface of cerebral cortex, along with the full cortical ROI parcellation of the SpeechLabel atlas described in Cai et al. (2014). Target cluster 4 is displayed on a transparent 3D rendering of the cerebellum (posterior view). The bottom portion of the figure shows the connectivity effect sizes for each connection in the normal and rhythm conditions, separately for each group. Error bars indicate 90% confidence intervals. N = normal, R = rhythm, L = left, R = right, vPMC = ventral premotor cortex, Cbm = cerebellum.

### Discussion

This study aimed to characterize the changes in functional activation and connectivity that occur when adults time their speech to an external metronomic beat and how these changes differ in AWS compared to ANS. Extending previous work, this paradigm was novel in that the metronome was paced at the typical rate of English speech. The rate and rhythmicity of paced speech by AWS was also similar to that of ANS. Consistent with prior literature, AWS produced significantly fewer disfluencies during externally-paced speech than during normal, internally-paced speech (Figure 3). In addition, while ANS exhibited greater activation during rhythmic speech than normal speech in left hemisphere auditory, premotor, and sensory association areas, as well as right hemisphere premotor cortex, AWS did not exhibit any significant differences between the conditions. AWS also had greater functional connectivity during rhythmic speech than normal speech between bilateral inferior cerebellum and orbitofrontal cortex and among all cerebellar speech regions. Finally, functional connections between right cerebellum and medial sensorimotor cortex and between both left vPMC and right putamen and right prefrontal cortex were significantly modulated by group and condition. The following sections discuss these results in relation to prior behavioral and neuroimaging literature.

#### A Compensatory Role for the Cerebellum in AWS

The role of the cerebellum for mediating speech timing is well-known (see Ackermann, 2008 for a review), and damage to this structure can lead to “scanning speech,” where syllables are evenly paced (Duffy, 2013). Previous work posits that when the basal-ganglia-SMA “internal” timing system is impaired in AWS, the cerebellum, along with lateral cortical premotor structures, forms part of an “external” timing system that is recruited (Alm, 2004; Etchell et al., 2014). In support of this, numerous fMRI and PET studies demonstrate cerebellar overactivation and hyper-connectivity during normal speech production in AWS (e.g., Brown et al., 2005; Chang et al., 2009; Ingham et al., 2012; Lu, Peng, et al., 2010; Lu et al., 2012; Watkins et al., 2007) that is reduced following therapy (De Nil et al., 2001; Lu et al., 2012; Neumann et al., 2003; Toyomura et al., 2015), a potential indication of an organic attempt at compensation. In the present study, the increased connectivity among speech-related regions of the cerebellum along with increased fluency during the rhythm condition may thus reflect similar neural processes.

It should be noted that this functional connectivity likely does not necessarily reflect direct structural connectivity between a seed and target region. Except in the case of connectivity between the cerebellar cortex and the dentate nucleus, which are structurally connected, viral tracing studies have found that each part of the cerebellar cortex forms closed-loop circuits with areas of cerebral cortex (Strick et al., 2009), meaning that different parts of cerebellar cortex do not communicate directly. Nonetheless, as suggested by (Bernard et al., 2013), we interpret the result of increased “within-cerebellar” connectivity as reflecting an increase in synchrony among multiple cerebro-cerebellar loops. Thus, in AWS, areas of cerebral cortex may simultaneously impinge on distinct areas of cerebellum to utilize the cerebellum’s temporal processing capabilities to ensure accurate speech timing during the *rhythm* condition.

The orbitofrontal cortex has also been shown to play a role in increasing fluency. Previous work on the OFC in AWS have shown greater OFC activation during speech in more fluent speakers (Kell et al., 2009), greater OFC activity following therapy for AWS (Kell et al., 2009), and increased activation in adults who spontaneously recovered from stuttering during adulthood in the left OFC compared to both persistent AWS and controls (Kell et al., 2009). The current study did not show greater activation in the OFC, but did show increased connectivity with the cerebellum during rhythmic speech. Previous studies have also found a relationship between increased functional connectivity between the cerebellum and the OFC and decreased stuttering severity in AWS (Sitek et al., 2016) and in adults who spontaneously recovered from stuttering during adulthood compared to ANS (Kell et al., 2018). Thus, increased connectivity between the cerebellum and OFC may underpin successful long-term compensatory behavior (i.e. fluency), which is induced by the rhythm condition in the current study.

There were also cerebellar connections that showed significant interactions between groups and conditions whereby the rhythm condition had the opposite effect on connectivity in the two groups. The AWS group had increased connectivity between right cerebellar lobule V and another cluster in posterior cerebellum, while the ANS had decreased connectivity. This increase in the AWS supports the earlier argument that increased connectivity within the cerebellum may reflect a compensatory mechanism. The AWS group also had decreased connectivity between the right cerebellum lobule V and left medial sensorimotor cortex and SMA, while the ANS group had increased connectivity between these areas. This may reflect that AWS have positive connectivity between the cerebellum (“external”) and medial premotor (“internal”) areas in the normal condition to compensate for the impaired “internal” basal-ganglia timing system. This connection is decreased in the rhythm condition because the AWS no longer attempt to use the medial structures. Conversely, ANS may have increased connectivity between these regions in the rhythm condition because the internal system is both working properly and is being used to a greater extent as seen in the task activation results. Together, all of these results support the theory that in AWS, an “external” timing system mediated by the cerebellum plays an increased role in speech production during externally-timed speech and can lead to increased fluency.

#### Increased Prefrontal Mediation During Rhythmic Speech

The AWS group also had decreased functional connections between right aMFG and both right vPMC and right putamen during the rhythm condition, whereas the ANS had increased connectivity. The right aMFG, a portion of dorsolateral prefrontal cortex, has been previously implicated in high-level cognitive tasks that require holding multiple pieces of information in memory (Barbey et al., 2013; Wager & Smith, 2003), including reframing emotional situations (Falquez et al., 2014; Ochsner et al., 2012). In the context of stuttering, it is well known that people who stutter will often monitor their upcoming speech in order to anticipate and potentially correct disfluencies (Garcia-Barrera & Davidow, 2015; Jackson et al., 2015). However, after noticing how speaking along with a metronome improves fluency, they may be less likely to continuously monitor their speech to the same extent. Therefore, decreased connectivity between right aMFG and left vPMC, an area hypothesized to encode speech motor programs (Guenther, 2016), in the rhythm condition may reflect this decreased monitoring of upcoming speech. As connections between lateral prefrontal cortex and basal ganglia structures mediate attention-shifting (Morris et al., 2016), decreased monitoring may also lead to decreased connectivity between right aMFG and right putamen. ANS, on the other hand, may exhibit an increase in connectivity between these regions because there is no fluency advantage to rhythmic speech and may require more monitoring to speak rhythmically. This conscious shift in attention may be mediated by increased connectivity between right putamen and right aMFG in ANS (Morris et al., 2016). Thus, the interaction found between group and condition in functional connections between speech planning and sequencing areas and right aMFG may be reflective of different changes in attentional demands between groups.

#### Changes in Activation due to Rhythmically-Timed Speech

Comparing neural activation between rhythmic and normal speech showed that ANS had greater activation during rhythmic speech than normal speech in left hemisphere auditory, premotor, and sensory association areas, as well as right hemisphere ventral premotor cortex. Activation in left auditory associative cortex (PT, PP) and right ventral premotor cortex (vPMC) may be related to increased reliance on auditory feedback control during this novel speech condition. Previous studies have shown that auditory feedback errors lead to increased activation in posterior auditory areas (Hashimoto & Sakai, 2003; Parkinson et al., 2012; Takaso et al., 2010; Tourville et al., 2008), and greater activation in right vPMC is thought to generate corrective responses to sensory errors in response to this altered sensory feedback (Golfinopoulos et al., 2011; Hashimoto & Sakai, 2003; Tourville et al., 2008). Alternatively, left PT has been described as an auditory-motor interface (Hickok et al., 2003); therefore increased activation in left PT may be indicative of the need to hold the rhythmic auditory stimulus in working memory and translate it into a motoric response in the rhythm condition of the current study. This is supported by increased activity (found in the exploratory analysis) in left anterior insula and superior parietal lobule, additional regions commonly recruited in working memory tasks (Rottschy et al., 2012).

There was also increased activation during rhythmic speech in areas thought to be involved in speech planning and sequencing (left SMA, pre-SMA, caudate and VA; Bohland et al., 2010; Civier et al., 2013; Guenther, 2016), articulatory planning of complex sequences (left aINS; (Ackermann & Riecker, 2010; Bohland & Guenther, 2006; Shuster & Lemieux, 2005), producing complex motor sequences (left SPL; Haslinger et al., 2002; Heim et al., 2012), producing untrained sequences (left SPL; Jenkins et al., 1994; Segawa et al., 2015), and attending to stimulus timing (left SPL; Coull, 2004). The rhythm condition requires participants to produce speech in an unfamiliar way. This change in their speech production results in speech becoming less automatic, and may require greater recruitment in these areas for timing the sequence of syllables (Alario et al., 2006; Bohland & Guenther, 2006; Schubotz & von Cramon, 2001). Bengtsson et al. (2004, 2005) found that for both finger tapping and simple repetition of “pa,” more complex timing led to increased activation in SMA and preSMA compared to simple patterns. The increased need to implement a timing pattern recruited these same structures that mediate temporal sequencing.

Unlike previous studies (Braun et al., 1997; Stager et al., 2003; Toyomura et al., 2011, 2015), AWS did not exhibit significantly increased activation in the *rhythm* condition compared to the *normal* condition. The most consistent finding from these studies was that both groups showed increased activation in bilateral auditory regions during rhythmic speech and that AWS showed greater increases in the basal ganglia. In the present study, the lack of clear between-condition effects within the AWS or between the AWS and ANS group may be due to more individual variability for AWS than ANS for this contrast. Future work is needed to determine whether this within-group variability is driving the null findings in the AWS group. Furthermore, Toyomura et al. (2011) found that while areas of the basal ganglia, left precentral gyrus, left SMA, left IFG, and left insula were less active in AWS during normal speech, activity in these areas increased to the level of ANS during rhythmic speech. These results suggested that rhythmic speech had a “normalizing” effect on activity in these regions, which differs with the present results.

There are methodological differences between the current work and similar studies that also could have impacted the results. In the current study, the rhythmic stimulus was presented prior to speaking regardless of the condition, unlike previous work in which the participant heard the stimulus while speaking and only during the rhythmic condition (Toyomura et al. 2011). Thus, group effects reported by Toyomura and colleagues (2011) may reflect differences in processing the auditory pacing stimulus in addition to differences in speech motor processes. Second, our study sought to examine the rhythm effect when speech was produced at a conversational speaking rate. Previous studies used a metronome set at 92 - 100 beats per minute, considerably slower than the mean conversational rate in English (228 - 372 syllables per minute; Davidow, 2014; Pellegrino et al., 2011) and the rate observed in our study (approximately 207 syllables per minute). While Toyomura et al. (2011, 2015) instructed participants to speak at a similar rate during the normal condition (when previous studies had not), the slower tempo overall may have led to increased auditory feedback processing. This could have modified the mechanisms by which ANS and AWS controlled their speech timing. Finally, only one of the previous studies accounted for disfluencies during the task in their imaging analysis (Stager et al., 2003), despite significant correlations with brain activation (Braun et al., 1997). However, given the small number of disfluencies in this and previous studies, this effect may have had a limited impact on the results.

#### Correlation Between Activation and Severity

The primary analysis found significant positive correlations between Disfluency Rate and activation in the *Normal-Baseline* contrast in left VA thalamus and bilateral VL thalamus. These nuclei are part of both the cortico-cerebellar and cortico-basal ganglia motor loops, and are structurally connected with premotor and primary motor areas (Barbas et al., 2013). As relays between subcortical structures and the cortex, increased activation for participants with a higher disfluency rate during the task may reflect greater reliance upon these modulatory pathways during speech. It is also worth noting that with an exploratory threshold (p < 0.05, uncorrected), some ROIs follow similar patterns to previous literature. Higher SSI-Mod scores were associated with weaker activation in left auditory areas. This correlation has been shown before (Fox et al., 2000) and there are numerous reports of atypical activity and/or morphology in left auditory cortex in AWS (e.g. Belyk et al., 2015; Chang et al., 2009; De Nil et al., 2000, 2008; Fox et al., 1996; Stager et al., 2003; Van Borsel et al., 2003). Similarly, the propensity to stutter during the task, measured by Disfluency Rate, is associated with greater cortical activation in largely right hemisphere regions, and bilateral subcortical activation at uncorrected thresholds. The right-lateralized cortical associations in the present study may reflect increased compensatory activity in AWS (as in Braun et al., 1997; Cai et al., 2014; Kell et al., 2009; Preibisch et al., 2003; Salmelin et al., 2000). This is supported by the fact that fluency-inducing therapy lead to more left-lateralized activation (De Nil et al., 2003; Neumann et al., 2003, 2005), similar to that of neurotypical speakers. It should be noted that due to the low number of disfluencies exhibited during the task, determining a clear relationship between stuttering severity and activation may not have been possible.

#### Limitations

Despite training in a prior session and feedback immediately prior to scanning, participants’ *rhythmic* speech productions were significantly slower than their *normal* speech productions. Since rate reduction is another method that reduces disfluencies in PWS (Andrews et al., 1982), this potentially could have led to the changes in both fluency and brain activation found herein. This same issue was reported in one previous neuroimaging study of the metronome-timed speech effect (Toyomura et al., 2011). As previously mentioned, the effect on rhythmic speech on fluency occurs even at high speaking rates (Davidow, 2014). Additionally, studies examining the effect of speaking rate on brain activation have found positive correlations with activation in sensorimotor cortex, SMA, insula, thalamus, and cerebellum (Fox et al., 2000; Riecker et al., 2006). This is the opposite effect of what we would expect given that increased activation was found (in ANS) during the slower *rhythmic* condition. While not conclusive, this evidence mitigates the concern that a decreased speaking rate accounted for neural changes found in this study.

In addition, the current results are not consistent with a recent meta-analysis examining activation differences between AWS and ANS (Belyk et al., 2015, 2017) which found that AWS consistently had overactivation in right hemisphere cortical structures, and underactivation in left hemisphere structures, especially in motor and premotor areas. However, the present study’s exploratory analysis suggested that AWS had decreased activation in left frontal operculum during the rhythmic condition as compared to the ANS group. Previous work has shown gray matter and white matter anomalies in and near left IFG (Beal et al., 2013, 2015; Chang et al., 2008, 2011; Kell et al., 2009; Lu et al., 2012), which may be related to this under-activation. Based on the exploratory nature of these findings, future work as well as meta-analytic testing is needed to determine whether these are true population differences.

## Conclusion

In this study, we examined brain activation patterns that co-occur with the introduction of an external pacing stimulus. We found that AWS showed an overall decrease in disfluencies during this condition, as well as functional connectivity changes between the cerebellum, prefrontal cortex, and other regions of the speech production network. Involvement of these structures suggests that rhythmic speech activates compensatory timing systems and potentially enhances top-down feedback control and attentional systems. This study provides greater insight into the network of brain areas that either support (or respond to) fluency in relation to the rhythm effect and its correspondence to longer-term fluency provided through natural compensation or therapy. It is our hope that in conjunction with the large body of work already published on fluency-enhancing techniques and future studies with more focused analyses, the field will come to a better understanding of the pathophysiology of stuttering and fluency, and that this information will be used to provide more targeted treatments and, ultimately, improve quality of life for those who stutter.

## Acknowledgements

The research reported here was supported by grants from the National Institute of Health (R01 DC007683, T32 DC013017, P41 RR14075) and the National Science Foundation (NSF 1625552). We are grateful to Diane Constantino, Barbara Holland, Matthias Heyne, Megan Thompson, Elaine Kearney, Julianne Leber, and Erin Archibald for assistance with subject recruitment and data collection and Ina Jessen, Mona Tong and Brittany Steinfeld for their help with behavioral data analysis. This work benefited from helpful discussions with, or comments from, other members of the Boston University Speech Lab.

## Appendix Stimulus sentences used in the present experiment

1. Rice is often served in round bowls.
2. The juice of lemons makes fine punch.
3. The boy was there when the sun rose.
4. Her purse was full of useless trash.
5. Hoist the load to your left shoulder.
6. The young girl gave no clear response.
7. Sickness kept him home the third week.
8. Lift the square stone over the fence.
9. The friendly gang left the drug store.
10. The lease ran out in sixteen weeks.
11. The steady bat gave birth to pups.
12. There are more than two factors here.
13. The lawyer tried to lose his case.
14. The term ended late June that year.
15. The pipe began to rust while new.
16. Act on these orders with great speed.

## References

Ackermann, H. (2008). Cerebellar contributions to speech production and speech perception: Psycholinguistic and neurobiological perspectives. Trends in Neurosciences, 31(6), 265–272. https://doi.org/10.1016/j.tins.2008.02.011

Ackermann, H., & Riecker, A. (2010). The contributions) of the insula to speech production: A review of the clinical and functional imaging literature. Brain Structure and Function, 214(5–6), 419–433. https://doi.org/10.1007/s00429-010-0257-x

Alario, F.-X., Chainay, H., Lehericy, S., & Cohen, L. (2006). The role of the supplementary motor area (SMA) in word production. Brain Research, 1076(1), 129–143. https://doi.org/10.1016/j.brainres.2005.11.104

Alm, P. A. (2004). Stuttering and the basal ganglia circuits: A critical review of possible relations. Journal of Communication Disorders, 37(4), 325–369. https://doi.org/10.1016/j.jcomdis.2004.03.001

Andersson, J. L. R., Hutton, C., Ashburner, J., Turner, R., & Friston, K. (2001). Modeling Geometric Deformations in EPI Time Series. NeuroImage, 13(5), 903–919. https://doi.org/10.1006/nimg.2001.0746

Andrews, G., Howie, P. M., Dozsa, M., & Guitar, B. E. (1982). Stuttering: Speech pattern characteristics under fluency-inducing conditions. Journal of Speech and Hearing Research, 25(2), 208–216.

Ashburner, J., & Friston, K. J. (2005). Unified segmentation. NeuroImage, 26(3), 839–851. https://doi.org/10.1016/j.neuroimage.2005.02.018

Azrin, N., Jones, R. J., & Flye, B. (1968). A synchronization effect and its application to stuttering by a portable apparatus1. Journal of Applied Behavior Analysis, 1(4), 283–295. https://doi.org/10.1901/jaba.1968.1-283

Barbas, H., García-Cabezas, M. Á., & Zikopoulos, B. (2013). Frontal-thalamic circuits associated with language. Brain and Language, 126(1), 49–61. https://doi.org/10.1016/j.bandl.2012.10.001

Barber, V. (1940). Studies in the Psychology of Stuttering, XVI: Rhythm as a Distraction in Stuttering. Journal of Speech Disorders, 5(1), 29–42. https://doi.org/10.1044/jshd.0501.29

Barbey, A. K., Koenigs, M., & Grafman, J. (2013). Dorsolateral prefrontal contributions to human working memory. Cortex, 49(5), 1195–1205. https://doi.org/10.1016/j.cortex.2012.05.022

Beal, D. S., Gracco, V. L., Brettschneider, J., Kroll, R. M., & De Nil, L. F. (2013). A voxel-based morphometry (VBM) analysis of regional grey and white matter volume abnormalities within the speech production network of children who stutter. Cortex, 49(8), 2151–2161. https://doi.org/10.1016/j.cortex.2012.08.013

Beal, D. S., Lerch, J. P., Cameron, B., Henderson, R., Gracco, V. L., & De Nil, L. F. (2015). The trajectory of gray matter development in Brocaâ€^TM^s area is abnormal in people who stutter. Frontiers in Human Neuroscience, 9. https://doi.org/10.3389/fnhum.2015.00089

Behzadi, Y., Restom, K., Liau, J., & Liu, T. T. (2007). A component based noise correction method (CompCor) for BOLD and perfusion based fMRI. NeuroImage, 37(1), 90–101. https://doi.org/10.1016/j.neuroimage.2007.04.042

Belin, P., Zatorre, R. J., Hoge, R., Evans, A. C., & Pike, B. (1999). Event-Related fMRI of the Auditory Cortex. NeuroImage, 10(4), 417–429. https://doi.org/10.1006/nimg.1999.0480

Belyk, M., Kraft, S. J., & Brown, S. (2015). Stuttering as a trait or state—An ALE meta-analysis of neuroimaging studies. European Journal of Neuroscience, 41(2), 275–284. https://doi.org/10.1111/ejn.12765

Belyk, M., Kraft, S. J., & Brown, S. (2017). Stuttering as a trait or a state revisited: Motor system involvement in persistent developmental stuttering. European Journal of Neuroscience, 45(4), 622–624. https://doi.org/10.1111/ejn.13512

Bengtsson, S. L., Ehrsson, H. H., Forssberg, H., & Ullen, F. (2004). Dissociating brain regions controlling the temporal and ordinal structure of learned movement sequences. European Journal of Neuroscience, 19(9), 2591–2602. https://doi.org/10.1111/j.0953-816X.2004.03269.x

Bengtsson, S. L., Ehrsson, H. H., Forssberg, H., & Ullén, F. (2005). Effector-independent voluntary timing: Behavioural and neuroimaging evidence. European Journal of Neuroscience, 22(12), 3255–3265. https://doi.org/10.1111/j.1460-9568.2005.04517.x

Benjamini, Y., & Hochberg, Y. (1995). Controlling the False Discovery Rate: A Practical and Powerful Approach to Multiple Testing. Journal of the Royal Statistical Society: Series B (Methodological), 57(1), 289–300. https://doi.org/10.1111/j.2517-6161.1995.tb02031.x

Bernard, J. A., Peltier, S. J., Wiggins, J. L., Jaeggi, S. M., Buschkuehl, M., Fling, B. W., Kwak, Y., Jonides, J., Monk, C. S., & Seidler, R. D. (2013). Disrupted cortico-cerebellar connectivity in older adults. NeuroImage, 83, 103–119. https://doi.org/10.1016/j.neuroimage.2013.06.042

Bohland, J. W., Bullock, D., & Guenther, F. H. (2010). Neural representations and mechanisms for the performance of simple speech sequences. Journal of Cognitive Neuroscience, 22(7), 1504–1529.

Bohland, J. W., & Guenther, F. H. (2006). An fMRI investigation of syllable sequence production. NeuroImage, 32(2), 821–841. https://doi.org/10.1016/j.neuroimage.2006.04.173

Braun, A. R., Varga, M., Stager, S., Schulz, G., Selbie, S., Maisog, J. M., Carson, R. E., & Ludlow, C. L. (1997). Altered patterns of cerebral activity during speech and language production in developmental stuttering. An H2 (15) O positron emission tomography study. Brain, 120(5), 761–784.

Brown, S., Ingham, R. J., Ingham, J. C., Laird, A. R., & Fox, P. T. (2005). Stuttered and fluent speech production: An ALE meta-analysis of functional neuroimaging studies. Human Brain Mapping, 25(1), 105–117. https://doi.org/10.1002/hbm.20140

Cai, S., Tourville, J. A., Beal, D. S., Perkell, J. S., Guenther, F. H., & Ghosh, S. S. (2014). Diffusion imaging of cerebral white matter in persons who stutter: Evidence for network-level anomalies. Frontiers in Human Neuroscience, 8. https://doi.org/10.3389/fnhum.2014.00054

Chang, S.-E., Erickson, K. I., Ambrose, N. G., Hasegawa-Johnson, M. A., & Ludlow, C. L. (2008). Brain anatomy differences in childhood stuttering. NeuroImage, 39(3), 1333–1344. https://doi.org/10.1016/j.neuroimage.2007.09.067

Chang, S.-E., Horwitz, B., Ostuni, J., Reynolds, R., & Ludlow, C. L. (2011). Evidence of Left Inferior Frontal–Premotor Structural and Functional Connectivity Deficits in Adults Who Stutter. Cerebral Cortex, 21(11), 2507–2518. https://doi.org/10.1093/cercor/bhr028

Chang, S.-E., Kenney, M. K., Loucks, T. M. J., & Ludlow, C. L. (2009). Brain activation abnormalities during speech and non-speech in stuttering speakers. NeuroImage, 46(1), 201–212. https://doi.org/10.1016/j.neuroimage.2009.01.066

Chang, S.-E., & Zhu, D. C. (2013). Neural network connectivity differences in children who stutter. Brain, 136(12), 3709–3726. https://doi.org/10.1093/brain/awt275

Chauvigné, L. A. S., Gitau, K. M., & Brown, S. (2014). The neural basis of audiomotor entrainment: An ALE meta-analysis. Frontiers in Human Neuroscience, 8. https://doi.org/10.3389/fnhum.2014.00776

Civier, O., Bullock, D., Max, L., & Guenther, F. H. (2013). Computational modeling of stuttering caused by impairments in a basal ganglia thalamo-cortical circuit involved in syllable selection and initiation. Brain and Language, 126(3), 263–278. https://doi.org/10.1016/j.bandl.2013.05.016

Coull, J. T. (2004). fMRI studies of temporal attention: Allocating attention within, or towards, time. Cognitive Brain Research, 21(2), 216–226. https://doi.org/10.1016/j.cogbrainres.2004.02.011

Craig, A., Blumgart, E., & Tran, Y. (2009). The impact of stuttering on the quality of life in adults who stutter. Journal of Fluency Disorders, 34(2), 61–71. https://doi.org/10.1016/j.jfludis.2009.05.002

Craig, A., & Tran, Y. (2006). Fear of speaking: Chronic anxiety and stammering. Advances in Psychiatric Treatment, 12(1), 63–68. https://doi.org/10.1192/apt.12.1.63

Craig, A., & Tran, Y. (2014). Trait and social anxiety in adults with chronic stuttering: Conclusions following meta-analysis. Journal of Fluency Disorders, 40, 35–43. https://doi.org/10.1016/j.jfludis.2014.01.001

Davidow, J. H. (2014). Systematic studies of modified vocalization: The effect of speech rate on speech production measures during metronome-paced speech in persons who stutter: Speech rate and speech production measures during metronome-paced speech in PWS. International Journal of Language & Communication Disorders, 49(1), 100–112. https://doi.org/10.1111/1460-6984.12050

De Nil, L. F., Kroll, R. M., & Houle, S. (2001). Functional neuroimaging of cerebellar activation during single word reading and verb generation in stuttering and nonstuttering adults. Neuroscience Letters, 302(2–3), 77–80. https://doi.org/10.1016/s0304-3940(01)01671-8

De Nil, Luc F., Beal, D. S., Lafaille, S. J., Kroll, R. M., Crawley, A. P., & Gracco, V. L. (2008). The effects of simulated stuttering and prolonged speech on the neural activation patterns of stuttering and nonstuttering adults. Brain and Language, 107(2), 114–123. https://doi.org/10.1016/j.bandl.2008.07.003

De Nil, Luc F., Kroll, R. M., Kapur, S., & Houle, S. (2000). A Positron Emission Tomography Study of Silent and Oral Single Word Reading in Stuttering and Nonstuttering Adults. Journal of Speech, Language, and Hearing Research, 43(4), 1038–1053. https://doi.org/10.1044/jslhr.4304.1038

De Nil, Luc F, Kroll, R. M., Lafaille, S. J., & Houle, S. (2003). A positron emission tomography study of short- and long-term treatment effects on functional brain activation in adults who stutter. Journal of Fluency Disorders, 28(4), 357–380. https://doi.org/10.1016/j.jfludis.2003.07.002

Diedrichsen, J. (2006). A spatially unbiased atlas template of the human cerebellum. NeuroImage, 33(1), 127–138. https://doi.org/10.1016/j.neuroimage.2006.05.056

Diedrichsen, J., Balsters, J. H., Flavell, J., Cussans, E., & Ramnani, N. (2009). A probabilistic MR atlas of the human cerebellum. NeuroImage, 46(1), 39–46. https://doi.org/10.1016/j.neuroimage.2009.01.045

Diedrichsen, J., Maderwald, S., Küper, M., Thürling, M., Rabe, K., Gizewski, E. R., Ladd, M. E., & Timmann, D. (2011). Imaging the deep cerebellar nuclei: A probabilistic atlas and normalization procedure. NeuroImage, 54(3), 1786–1794. https://doi.org/10.1016/j.neuroimage.2010.10.035

Duffy, J. R. (2013). Motor speech disorders: Substrates, differential diagnosis, and management (Third edition). Elsevier.

Eden, G. F., Joseph, J. E., Brown, H. E., Brown, C. P., & Zeffiro, T. A. (1999). Utilizing hemodynamic delay and dispersion to detect fMRI signal change without auditory interference: The behavior interleaved gradients technique. Magnetic Resonance in Medicine, 41(1), 13–20. https://doi.org/10.1002/(SICI)1522-2594(199901)41:1<13::AID-MRM4>3.0.CO;2-T

Etchell, A. C., Civier, O., Ballard, K. J., & Sowman, P. F. (2018). A systematic literature review of neuroimaging research on developmental stuttering between 1995 and 2016. Journal of Fluency Disorders, 55, 6–45. https://doi.org/10.1016/j.jfludis.2017.03.007

Etchell, A. C., Johnson, B. W., & Sowman, P. F. (2014). Behavioral and multimodal neuroimaging evidence for a deficit in brain timing networks in stuttering: A hypothesis and theory. Frontiers in Human Neuroscience, 8. https://doi.org/10.3389/fnhum.2014.00467

Falquez, R., Couto, B., Ibanez, A., Freitag, M. T., Berger, M., Arens, E. A., Lang, S., & Barnow, S. (2014). Detaching from the negative by reappraisal: The role of right superior frontal gyrus (BA9/32). Frontiers in Behavioral Neuroscience, 8. https://doi.org/10.3389/fnbeh.2014.00165

Fischl, B., Sereno, M. I., & Dale, A. M. (1999). Cortical Surface-Based Analysis. NeuroImage, 9(2), 195–207. https://doi.org/10.1006/nimg.1998.0396

Fox, P. T., Ingham, R. J., Ingham, J. C., Hirsch, T. B., Downs, J. H., Martin, C., Jerabek, P., Glass, T., & Lancaster, J. L. (1996). A PET study of the neural systems of stuttering. Nature, 382(6587), 158–162. https://doi.org/10.1038/382158a0

Fox, P. T., Ingham, R. J., Ingham, J. C., Zamarripa, F., Xiong, J. H., & Lancaster, J. L. (2000). Brain correlates of stuttering and syllable production. A PET performance-correlation analysis. Brain: A Journal of Neurology, 123 (Pt 10), 1985–2004.

Garcia-Barrera, M. A., & Davidow, J. H. (2015). Anticipation in stuttering: A theoretical model of the nature of stutter prediction. Journal of Fluency Disorders, 44, 1–15. https://doi.org/10.1016/j.jfludis.2015.03.002

Garnett, E. O., Chow, H. M., Nieto-Castañón, A., Tourville, J. A., Guenther, F. H., & Chang, S. E. (2018). Anomalous morphology in left hemisphere motor and premotor cortex of children who stutter. Brain, 141(9), 2670–2684. https://doi.org/10.1093/brain/awy199

Giraud, A. (2008). Severity of dysfluency correlates with basal ganglia activity in persistent developmental stuttering. Brain and Language, 104(2), 190–199. https://doi.org/10.1016/j.bandl.2007.04.005

Golfinopoulos, E., Tourville, J. A., Bohland, J. W., Ghosh, S. S., Nieto-Castanon, A., & Guenther, F. H. (2011). FMRI investigation of unexpected somatosensory feedback perturbation during speech. NeuroImage, 55(3), 1324–1338. https://doi.org/10.1016/j.neuroimage.2010.12.065

Gracco, V. L., Tremblay, P., & Pike, B. (2005). Imaging speech production using fMRI. NeuroImage, 26(1), 294–301. https://doi.org/10.1016/j.neuroimage.2005.01.033

Guenther, F. H. (2016). Neural control of speech. MIT Press.

Guitar, B. (2014). Stuttering: An integrated approach to its nature and treatment (4th ed). Wolters Kluwer Health/Lippincott Williams & Wilkins.

Hall, D. A., Haggard, M. P., Akeroyd, M. A., Palmer, A. R., Summerfield, A. Q., Elliott, M. R., Gurney, E. M., & Bowtell, R. W. (1999). “Sparse” temporal sampling in auditory fMRI. Human Brain Mapping, 7(3), 213–223. https://doi.org/10.1002/(sici)1097-0193(1999)7:3<213::aid-hbm5>3.0.co;2-n

Hanna, R., & Morris, S. (1977). Stuttering, Speech Rate, and the Metronome Effect. Perceptual and Motor Skills, 44(2), 452–454. https://doi.org/10.2466/pms.1977.44.2.452

Hashimoto, Y., & Sakai, K. L. (2003). Brain activations during conscious self-monitoring of speech production with delayed auditory feedback: An fMRI study. Human Brain Mapping, 20(1), 22–28. https://doi.org/10.1002/hbm.10119

Haslinger, B., Erhard, P., Weilke, F., Ceballos-Baumann, A. O., Bartenstein, P., Gräfin von Einsiedel, H., Schwaiger, M., Conrad, B., & Boecker, H. (2002). The role of lateral premotor–cerebellar–parietal circuits in motor sequence control: A parametric fMRI study. Cognitive Brain Research, 13(2), 159–168. https://doi.org/10.1016/S0926-6410(01)00104-5

Heim, S., Amunts, K., Hensel, T., Grande, M., Huber, W., Binkofski, F., & Eickhoff, S. B. (2012). The Role of Human Parietal Area 7A as a Link between Sequencing in Hand Actions and in Overt Speech Production. Frontiers in Psychology, 3. https://doi.org/10.3389/fpsyg.2012.00534

Hickok, G., Buchsbaum, B., Humphries, C., & Muftuler, T. (2003). Auditory–Motor Interaction Revealed by fMRI: Speech, Music, and Working Memory in Area Spt. Journal of Cognitive Neuroscience, 15(5), 673–682. https://doi.org/10.1162/089892903322307393

Hutchinson, J. M., & Norris, G. M. (1977). The differential effect of three auditory stimuli on the frequency of stuttering behaviors. Journal of Fluency Disorders, 2(4), 283–293. https://doi.org/10.1016/0094-730X(77)90032-8

IEEE Recommended Practice for Speech Quality Measurements (No. 17; pp. 227–246). (1969). IEEE Transactions on Audio and Electroacoustics. https://doi.org/10.1109/IEEESTD.1969.7405210

Ingham, R. J., Fox, P. T., Costello Ingham, J., & Zamarripa, F. (2000). Is Overt Stuttered Speech a Prerequisite for the Neural Activations Associated with Chronic Developmental Stuttering? Brain and Language, 75(2), 163–194. https://doi.org/10.1006/brln.2000.2351

Ingham, R. J., Grafton, S. T., Bothe, A. K., & Ingham, J. C. (2012). Brain activity in adults who stutter: Similarities across speaking tasks and correlations with stuttering frequency and speaking rate. Brain and Language, 122(1), 11–24. https://doi.org/10.1016/j.bandl.2012.04.002

Jackson, E. S., Yaruss, J. S., Quesal, R. W., Terranova, V., & Whalen, D. H. (2015). Responses of adults who stutter to the anticipation of stuttering. Journal of Fluency Disorders, 45, 38–51. https://doi.org/10.1016/j.jfludis.2015.05.002

Jenkins, I., Brooks, D., Nixon, P., Frackowiak, R., & Passingham, R. (1994). Motor sequence learning: A study with positron emission tomography. The Journal of Neuroscience, 14(6), 3775–3790. https://doi.org/10.1523/JNEUROSCI.14-06-03775.1994

Kell, C. A., Neumann, K., Behrens, M., von Gudenberg, A. W., & Giraud, A.-L. (2018). Speaking-related changes in cortical functional connectivity associated with assisted and spontaneous recovery from developmental stuttering. Journal of Fluency Disorders, 55, 135–144. https://doi.org/10.1016/j.jfludis.2017.02.001

Kell, C. A., Neumann, K., von Kriegstein, K., Posenenske, C., von Gudenberg, A. W., Euler, H., & Giraud, A.-L. (2009). How the brain repairs stuttering. Brain, 132(10), 2747–2760. https://doi.org/10.1093/brain/awp185

Keuken, M. C., Bazin, P.-L., Crown, L., Hootsmans, J., Laufer, A., Müller-Axt, C., Sier, R., van der Putten, E. J., Schäfer, A., Turner, R., & Forstmann, B. U. (2014). Quantifying inter-individual anatomical variability in the subcortex using 7 T structural MRI. NeuroImage, 94, 40–46. https://doi.org/10.1016/j.neuroimage.2014.03.032

Krauth, A., Blanc, R., Poveda, A., Jeanmonod, D., Morel, A., & Székely, G. (2010). A mean three-dimensional atlas of the human thalamus: Generation from multiple histological data. NeuroImage, 49(3), 2053–2062. https://doi.org/10.1016/j.neuroimage.2009.10.042

Lee, A., & Kawahara, T. (2009, January). Recent Development of Open-Source Speech recognition Engine Julius. Asia-Pacific Signal and Information Processing Association, 2009 Annual Summit and Conference.

Lu, C., Chen, C., Ning, N., Ding, G., Guo, T., Peng, D., Yang, Y., Li, K., & Lin, C. (2010). The neural substrates for atypical planning and execution of word production in stuttering. Experimental Neurology, 221(1), 146–156. https://doi.org/10.1016/j.expneurol.2009.10.016

Lu, C., Chen, C., Peng, D., You, W., Zhang, X., Ding, G., Deng, X., Yan, Q., & Howell, P. (2012). Neural anomaly and reorganization in speakers who stutter: A short-term intervention study. Neurology, 79(7), 625–632. https://doi.org/10.1212/WNL.0b013e31826356d2

Lu, C., Ning, N., Peng, D., Ding, G., Li, K., Yang, Y., & Lin, C. (2009). The role of large-scale neural interactions for developmental stuttering. Neuroscience, 161(4), 1008–1026. https://doi.org/10.1016/j.neuroscience.2009.04.020

Lu, C., Peng, D., Chen, C., Ning, N., Ding, G., Li, K., Yang, Y., & Lin, C. (2010). Altered effective connectivity and anomalous anatomy in the basal ganglia-thalamocortical circuit of stuttering speakers. Cortex, 46(1), 49–67. https://doi.org/10.1016/j.cortex.2009.02.017

Max, L. (2004). Stuttering and internal models for sensorimotor control: A theoretical perspective to generate testable hypotheses. In B. Maassen, R. Kent, P. Hermann, & P. Van Lieshout (Eds.), Speech Motor Control: In Normal and Disordered Speech (pp. 357–387). Oxford University Press.

Max, L., Guenther, F. H., Gracco, V. L., Ghosh, S. S., & Wallace, M. E. (2004). Unstable or Insufficiently Activated Internal Models and Feedback-Biased Motor Control as Sources of Dysfluency: A Theoretical Model of Stuttering. Contemporary Issues in Communication Science and Disorders, 31(Spring), 105–122. https://doi.org/10.1044/cicsd_31_S_105

McLaren, D. G., Ries, M. L., Xu, G., & Johnson, S. C. (2012). A generalized form of context-dependent psychophysiological interactions (gPPI): A comparison to standard approaches. NeuroImage, 61(4), 1277–1286. https://doi.org/10.1016/j.neuroimage.2012.03.068

Morris, L. S., Kundu, P., Dowell, N., Mechelmans, D. J., Favre, P., Irvine, M. A., Robbins, T. W., Daw, N., Bullmore, E. T., Harrison, N. A., & Voon, V. (2016). Fronto-striatal organization: Defining functional and microstructural substrates of behavioural flexibility. Cortex, 74, 118–133. https://doi.org/10.1016/j.cortex.2015.11.004

Neumann, K., Euler, H. A., Gudenberg, A. W. von, Giraud, A.-L., Lanfermann, H., Gall, V., & Preibisch, C. (2003). The nature and treatment of stuttering as revealed by fMRI. Journal of Fluency Disorders, 28(4), 381–410. https://doi.org/10.1016/j.jfludis.2003.07.003

Neumann, K., Preibisch, C., Euler, H. A., Gudenberg, A. W. von, Lanfermann, H., Gall, V., & Giraud, A.-L. (2005). Cortical plasticity associated with stuttering therapy. Journal of Fluency Disorders, 30(1), 23–39. https://doi.org/10.1016/j.jfludis.2004.12.002

Nieto-Castañón, A. (2020). Handbook of functional connecitivity Magnetic resonance Imaging methods in CONN. Hilbert Press.

Ochsner, K. N., Silvers, J. A., & Buhle, J. T. (2012). Functional imaging studies of emotion regulation: A synthetic review and evolving model of the cognitive control of emotion: Functional imaging studies of emotion regulation. Annals of the New York Academy of Sciences, 1251(1), E1–E24. https://doi.org/10.1111/j.1749-6632.2012.06751.x

Oldfield, R. C. (1971). The assessment and analysis of handedness: The Edinburgh inventory. Neuropsychologia, 9(1), 97–113.

Parkinson, A. L., Flagmeier, S. G., Manes, J. L., Larson, C. R., Rogers, B., & Robin, D. A. (2012). Understanding the neural mechanisms involved in sensory control of voice production. NeuroImage, 61(1), 314–322. https://doi.org/10.1016/j.neuroimage.2012.02.068

Pellegrino, F., Coupé, C., & Marsico, E. (2011). Across-Language Perspective on Speech Information Rate. Language, 87(3), 539–558. https://doi.org/10.1353/lan.2011.0057

Pellegrino, F., Farinas, J., & Rouas, J.-L. (2004). Automatic Estimation of Speaking Rate in Multilingual Spontaneous Speech. Speech Prosody 2004, 517–520.

Preibisch, C., Neumann, K., Raab, P., Euler, H. A., von Gudenberg, A. W., Lanfermann, H., & Giraud, A.-L. (2003). Evidence for compensation for stuttering by the right frontal operculum. NeuroImage, 20(2), 1356–1364. https://doi.org/10.1016/S1053-8119(03)00376-8

Riecker, A., Kassubek, J., Gröschel, K., Grodd, W., & Ackermann, H. (2006). The cerebral control of speech tempo: Opposite relationship between speaking rate and BOLD signal changes at striatal and cerebellar structures. NeuroImage, 29(1), 46–53. https://doi.org/10.1016/j.neuroimage.2005.03.046

Riley, G. D. (2008). SSI-4, Stuttering severity intrument for children and adults (4th ed.). Pro Ed.

Rottschy, C., Langner, R., Dogan, I., Reetz, K., Laird, A. R., Schulz, J. B., Fox, P. T., & Eickhoff, S. B. (2012). Modelling neural correlates of working memory: A coordinate-based meta-analysis. NeuroImage, 60(1), 830–846. https://doi.org/10.1016/j.neuroimage.2011.11.050

Salmelin, R., Schnitzler, A., Schmitz, F., & Freund, H.-J. (2000). Single word reading in developmental stutterers and fluent speakers. Brain, 123(6), 1184–1202. https://doi.org/10.1093/brain/123.6.1184

Schubotz, R. I., & von Cramon, D. Y. (2001). Interval and Ordinal Properties of Sequences Are Associated with Distinct Premotor Areas. Cerebral Cortex, 11(3), 210–222. https://doi.org/10.1093/cercor/11.3.210

Segawa, J. A., Tourville, J. A., Beal, D. S., & Guenther, F. H. (2015). The Neural Correlates of Speech Motor Sequence Learning. Journal of Cognitive Neuroscience, 27(4), 819–831. https://doi.org/10.1162/jocn_a_00737

Shuster, L., & Lemieux, S. (2005). An fMRI investigation of covertly and overtly produced mono- and multisyllabic words. Brain and Language, 93(1), 20–31. https://doi.org/10.1016/j.bandl.2004.07.007

Sitek, K. R., Cai, S., Beal, D. S., Perkell, J. S., Guenther, F. H., & Ghosh, S. S. (2016). Decreased Cerebellar-Orbitofrontal Connectivity Correlates with Stuttering Severity: Whole-Brain Functional and Structural Connectivity Associations with Persistent Developmental Stuttering. Frontiers in Human Neuroscience, 10. https://doi.org/10.3389/fnhum.2016.00190

Stager, S. V., Denman, D. W., & Ludlow, C. L. (1997). Modifications in Aerodynamic Variables by Persons Who Stutter Under Fluency-Evoking Conditions. Journal of Speech, Language, and Hearing Research, 40(4), 832–847. https://doi.org/10.1044/jslhr.4004.832

Stager, S. V., Jeffries, K. J., & Braun, A. R. (2003). Common features of fluency-evoking conditions studied in stuttering subjects and controls: An PET study. Journal of Fluency Disorders, 28(4), 319–336. https://doi.org/10.1016/j.jfludis.2003.08.004

Strick, P. L., Dum, R. P., & Fiez, J. A. (2009). Cerebellum and Nonmotor Function. Annual

Review of Neuroscience, 32(1), 413–434. https://doi.org/10.1146/annurev.neuro.31.060407.125606

Takaso, H., Eisner, F., Wise, R. J. S., & Scott, S. K. (2010). The Effect of Delayed Auditory Feedback on Activity in the Temporal Lobe While Speaking: A Positron Emission Tomography Study. Journal of Speech, Language, and Hearing Research, 53(2), 226–236. https://doi.org/10.1044/1092-4388(2009/09-0009)

Teki, S., Grube, M., & Griffiths, T. D. (2012). A Unified Model of Time Perception Accounts for Duration-Based and Beat-Based Timing Mechanisms. Frontiers in Integrative Neuroscience, 5. https://doi.org/10.3389/fnint.2011.00090

Tourville, J. A., Reilly, K. J., & Guenther, F. H. (2008). Neural mechanisms underlying auditory feedback control of speech. NeuroImage, 39(3), 1429–1443. https://doi.org/10.1016/j.neuroimage.2007.09.054

Toyomura, A., Fujii, T., & Kuriki, S. (2011). Effect of external auditory pacing on the neural activity of stuttering speakers. NeuroImage, 57(4), 1507–1516. https://doi.org/10.1016/j.neuroimage.2011.05.039

Toyomura, A., Fujii, T., & Kuriki, S. (2015). Effect of an 8-week practice of externally triggered speech on basal ganglia activity of stuttering and fluent speakers. NeuroImage, 109, 458–468. https://doi.org/10.1016/j.neuroimage.2015.01.024

Van Borsel, J., Achten, E., Santens, P., Lahorte, P., & Voet, T. (2003). fMRI of developmental stuttering: A pilot study. Brain and Language, 85(3), 369–376. https://doi.org/10.1016/S0093-934X(02)00588-6

Wager, T. D., & Smith, E. E. (2003). Neuroimaging studies of working memory: Cognitive, Affective, & Behavioral Neuroscience, 3(4), 255–274. https://doi.org/10.3758/CABN.3.4.255

Watkins, K. E., Smith, S. M., Davis, S., & Howell, P. (2007). Structural and functional abnormalities of the motor system in developmental stuttering. Brain, 131(1), 50–59. https://doi.org/10.1093/brain/awm241

Whitfield-Gabrieli, S., & Nieto-Castanon, A. (2012). Conn: A Functional Connectivity Toolbox for Correlated and Anticorrelated Brain Networks. Brain Connectivity, 2(3), 125–141. https://doi.org/10.1089/brain.2012.0073

Wiener, M., Turkeltaub, P., & Coslett, H. B. (2010). The image of time: A voxel-wise meta-analysis. NeuroImage, 49(2), 1728–1740. https://doi.org/10.1016/j.neuroimage.2009.09.064

Yairi, E., & Ambrose, N. G. (1999). Early Childhood Stuttering I: Persistency and Recovery Rates. Journal of Speech Language and Hearing Research, 42(5), 1097. https://doi.org/10.1044/jslhr.4205.1097

Zeid, O., & Bullock, D. (2019). Moving in time: Simulating how neural circuits enable rhythmic enactment of planned sequences. Neural Networks, 120, 86–107. https://doi.org/10.1016/j.neunet.2019.08.006

