## Supplementary Figures and Tables for "The neural circuitry underlying the “rhythm effect” in stuttering"

**Supplementary Materials**

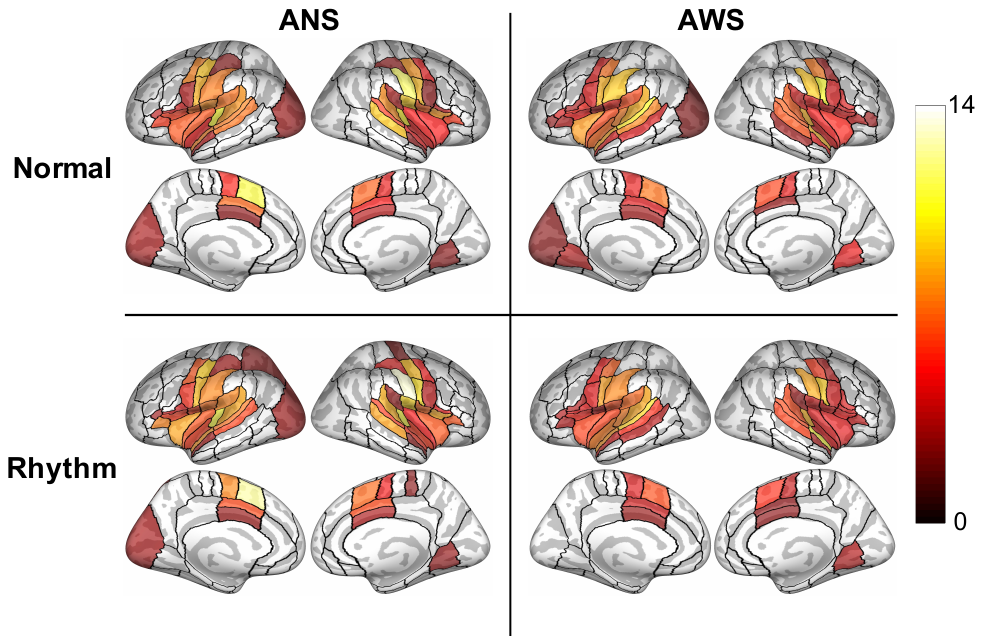
**Supplementary Figure 1.** Cortical regions-of-interest with significant positive activation plotted for each condition compared to baseline in each group using a one-sided t-test with a threshold of *p_FDR_* < 0.05. Results are presented on an inflated cortical surface, along with the full cortical ROI parcellation of the SpeechLabel atlas described in Cai et al. (2014). Color shading indicates t-values. ANS = adults who do not stutter, AWS = adults who stutter.

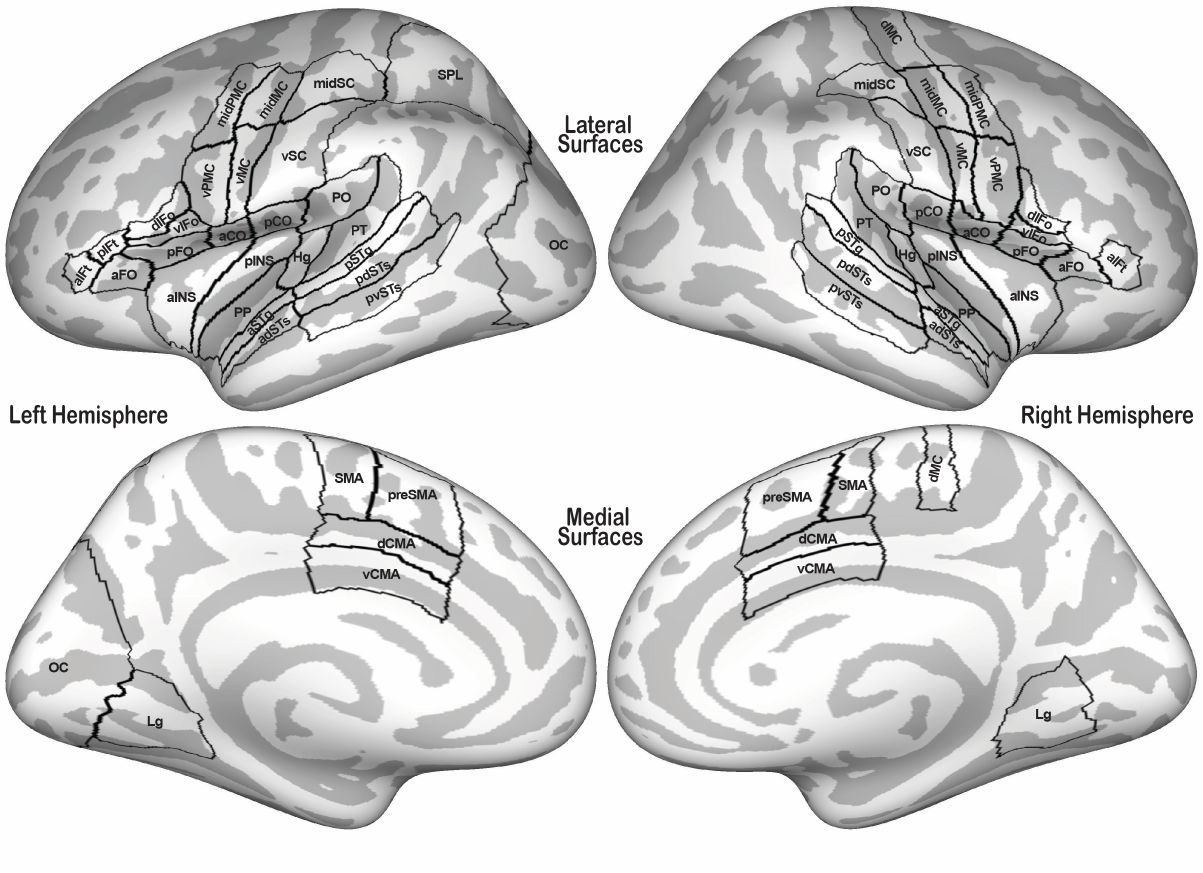
**Supplementary Figure 2.** Cortical regions-of-interest included in the exploratory analyses. aIFt = anterior inferior frontal gyrus pars triangularis, pIFt = posterior inferior temporal gyrus pars triangularis, aFO = anterior frontal operculum, pFO = posterior frontal operculum, dIFo = dorsal inferior frontal gyrus pars opercularis, vIFo = ventral inferior frontal gyrus pars opercularis, aINS = anterior insula, pINS = posterior insula, vPMC = ventral premotor cortex, midPMC = mid premotor cortex, vMC = ventral primary motor cortex, midMC = mid primary motor cortex, dMC = dorsal primary motor cortex, vSC = ventral somatosensory cortex, midSC = mid somatosensory cortex, aCO = anterior central operculum, pCO = posterior central operculum, PO = parietal operculum, PT = planum temporale, Hg = Heschl’s gyrus, PP = planum polare, aSTg = anterior superior temporal gyrus, pSTg = posterior superior temporal gyrus, adSTs = anterior dorsal superior temporal sulcus, pdSTs = posterior dorsal superior temporal sulcus, pvSTs = posterior ventral superior temporal sulcus, SPL = superior parietal lobule, OC = occipital cortex, Lg = lingual gyrus, SMA = supplementary motor area, preSMA = presupplementary motor area, dCMA = dorsal cingulate motor area, vCMA = ventral cingulate motor area.

**Supplementary Figure 3.** Subcortical regions-of-interest included in the exploratory analyses. L = left, R = right, Cbm = cerebellum, VA = ventroanterior thalamic nucleus, VL = ventrolateral thalamic nucleus, VPM = ventral posteromedial thalamic nucleus, MGN = medial geniculate nucleus of the thalamus, GPe = external portion of the globus pallidus, GPi = internal portion of the globus pallidus, STh = subthalamic nucleus, SN = substantia nigra.

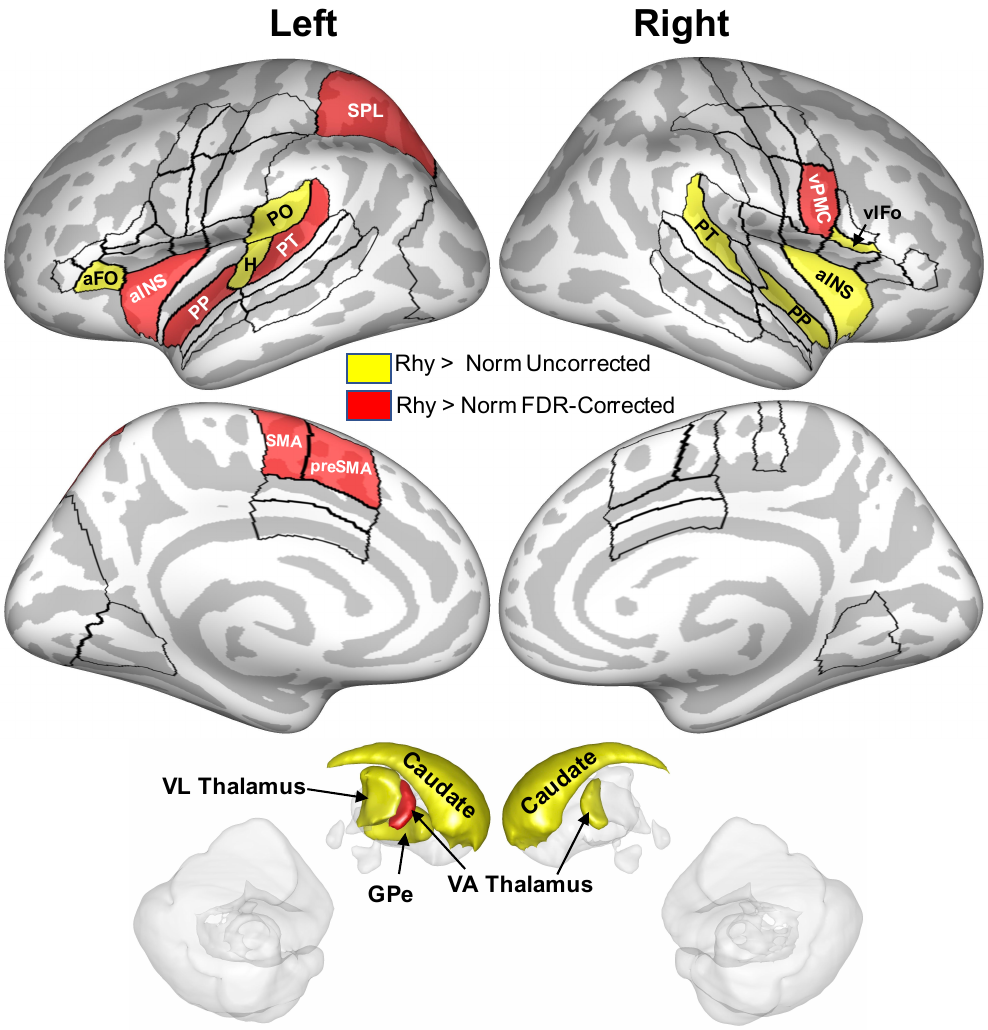

**Supplementary Figure 4.** Exploratory regions-of-interest (ROIs) significantly more active during the rhythmic condition than the normal condition for ANS in the exploratory analysis (*p* < 0.05) are highlighted in yellow and plotted on an inflated cortical surface. ROIs highlighted in red and labeled reached significance at a stricter threshold of *p_FDR_* < 0.05. Black outlines indicate cortical ROIs included in the exploratory analysis. FDR = false discovery rate, PT = planum temporale, SMA = supplementary motor area; preSMA = pre-supplementary motor area, SPL = superior parietal lobule, aINS = anterior insula, vPMC = ventral premotor cortex, VA = ventroanterior, PP = planum polare, vIFo = ventral inferior frontal gyrus pars opercularis, H = Heschl’s gyrus, aFO = anterior frontal operculum, VL = ventrolateral, GPe = external portion of the globus pallidus.

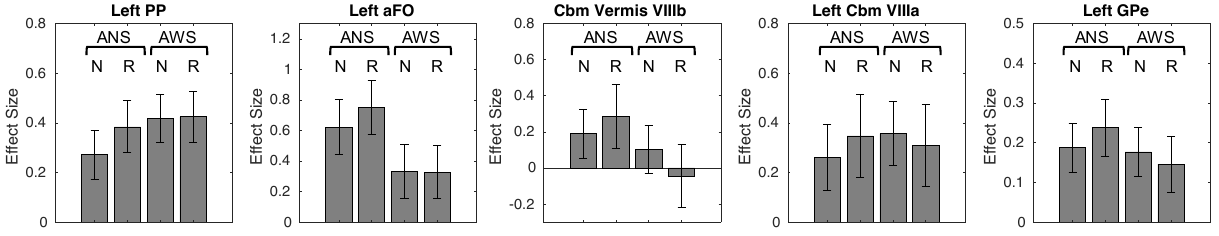

Supplementary Figure 5. Individual group and condition effects from the exploratory regions-of-interest that had a significant interaction between group and condition. PP = planum polare, aFO = anterior frontal operculum, Cbm = cerebellum, GPe = external portion of the globus pallidus, N = *Normal - Baseline* condition, R = *Rhythm - Baseline* condition, ANS = adults who do not stutter, AWS = adults who stutter. Error bars indicuate 90% confidence intervals.

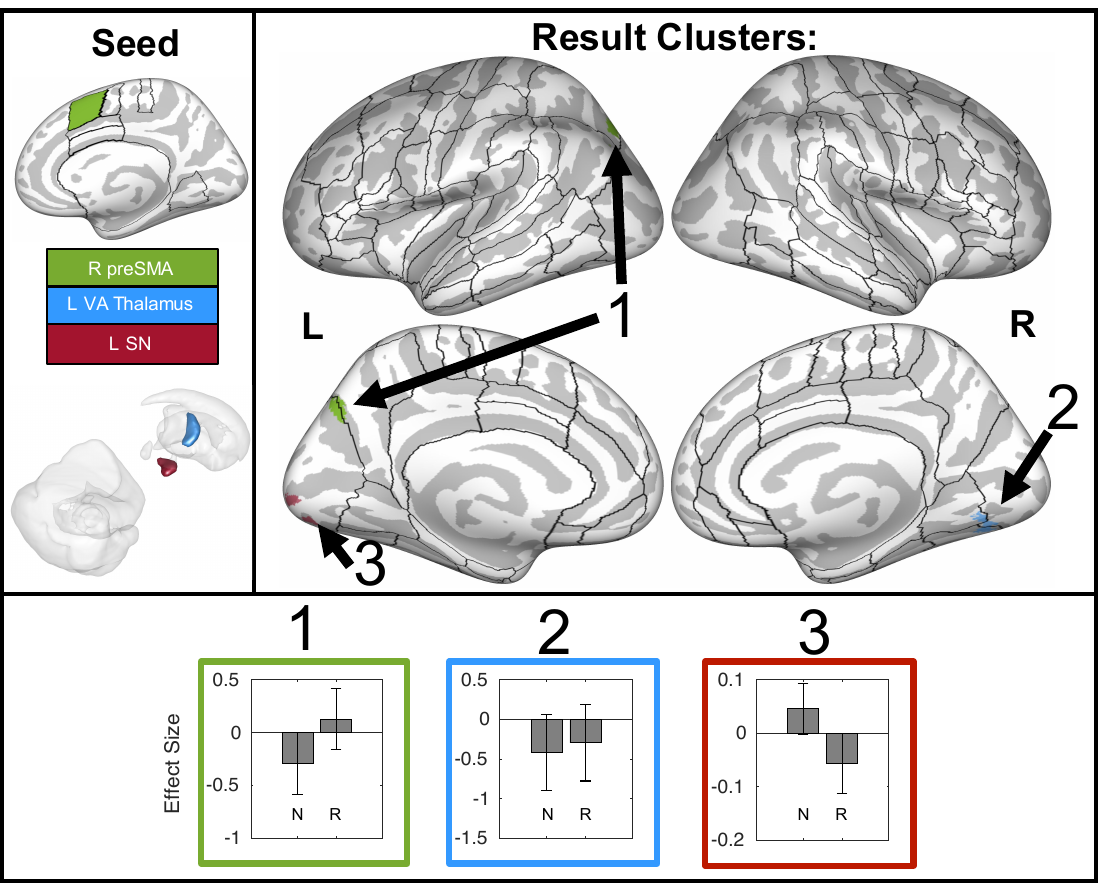

**Supplementary Figure 6.** A summary of functional connections that are significantly different between the *normal* and *rhythm* conditions in ANS. Seed regions for these connections are indicated in the upper left panel either on an inflated cortical surface (*top*; ROIs are as in Figure 2) or on a transparent 3D rendering of the left hemisphere subcortical structures viewed from the right (*bottom*). Colors in the rest of the figure refer back to these seed regions. Three target clusters (representing 3 distinct connections) are displayed in the upper right portion of the figure. These clusters are projected onto an inflated surface of cerebral cortex, along with the full cortical ROI parcellation of the SpeechLabel atlas described in Cai et al. (2014). The bottom portion of the figure shows the connectivity effect sizes in the *normal* and *rhythm* conditions for each connection. Error bars indicate 90% confidence intervals. N = normal, R = rhythm, L = left, R = right, preSMA = presupplementary motor area, VA = ventroanterior, SN = substantia nigra.

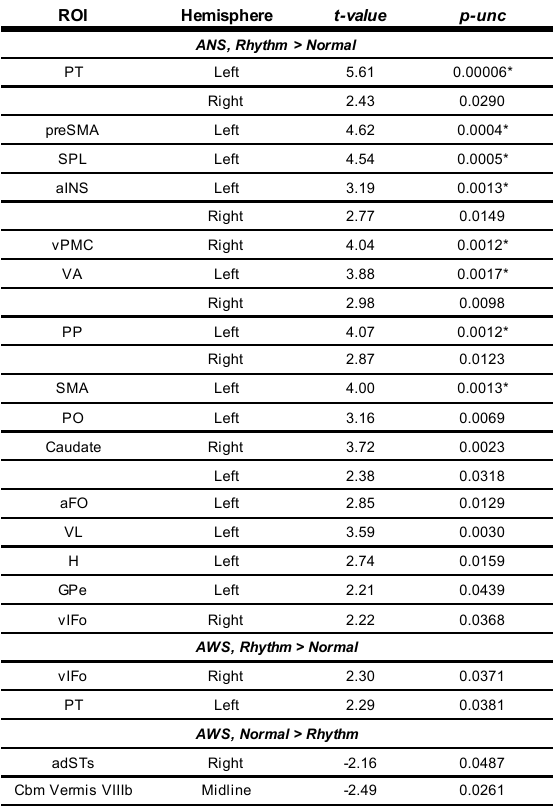

**Supplementary Table 1.** Exploratory regions-of-interest with activation differences between the *rhythm* and *normal* conditions for ANS and AWS (*p* < 0.05). * indicates regions that survive a significance threshold of *p_FDR_ <* 0.05 for their respective analyses, unc = uncorrested, PT = planum temporale, preSMA = presupplementary motor area, SPL = superior parietal lobule, aINS = anterior insula, vPMC = ventral premotor cortex, VA = ventroanterior thalamic nucleus, PP = planum polare; SMA = supplementary motor area, PO = parietal operculum, aFO = anterior frontal operculum, VL = ventrolateral thalamic nucleus, H = Heschl’s gyrus, GPe = external portion of the globus pallidus, vIFo = ventral inferior frontal gyrus pars opercularis, aSTg = anterior superior temporal gyrus, adSTs = anterior dorsal superior temporal sulcus, Cbm = cerebellum.

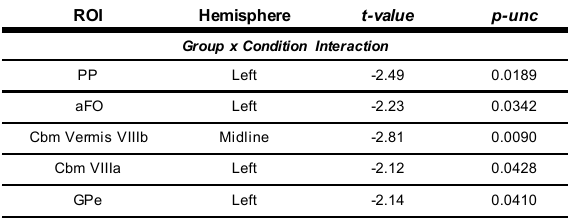

**Supplementary Table 2.** Exploratory regions-of-interest with significant task activation group x condition interactions (*p* < 0.05). unc = uncorrested, PP = planum polare, aFO = anterior frontal operculum, Cbm = cerebellum, GPe = external portion of globus pallidus.

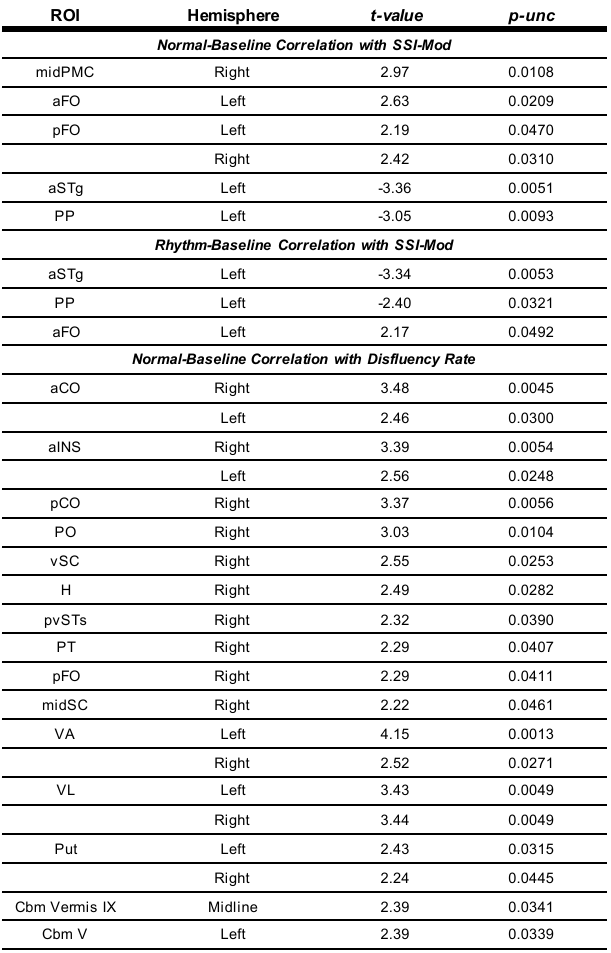

**Supplementary Table 3.** Exploratory regions-of-interest with significant correlations between severity measures and speech activation in AWS (*p* < 0.05). * indicates regions that survive a significance threshold of *p_FDR_ <* 0.05 for their respective analyses, unc = uncorrested, midPMC = mid premotor cortex, aSTg = anterior superior temporal gyrus, pdSTs = posterior dorsal superior temporal sulcus, aCO = anterior central operculum, pCO = posterior central operculum, PO = parietal operculum, aINS = anterior insula, vSC = ventral somatosensory cortex, H = Heschl’s gyrus, pvSTs = posterior ventral superior temporal sulcus, PT = planum temporale, pFO = posterior frontal operculum, midSC = mid somatosensory cortex, VA = ventroanterior thalamic nucleus, VL = ventrolateral thalamic nucleus, Put = putamen, Cbm = cerebellum.
